# Mitochondrial activity-driven hematopoietic stem cell fate and lympho-myeloid lineage choice is first established in the aorta-gonad-mesonephros

**DOI:** 10.1101/2024.08.25.609550

**Authors:** Aishwarya Prakash, Maneesha S. Inamdar

## Abstract

Mitochondrial metabolism determines bone marrow hematopoietic stem cell (HSC) heterogeneity and influences long-term repopulation potential. However, the origin of this heterogeneity and how it regulates HSC phenotype is unclear. We show that during the endothelial-to-hematopoietic transition (EHT) in the mouse embryo, dynamic changes in mitochondrial activity drive the production of mature HSCs with differing potencies. Lowering mitochondrial activity in the AGM by pharmacological or genetic means activates Wnt signaling to promote HSC expansion. Further, mitochondrial membrane potential (MMP) gives rise to functional heterogeneity in the definitive HSC pool. *In-vitro* and *in-vivo* functional assays, and single-cell transcriptomics on AGM HSCs showed that MMP^low^ HSCs are myeloid-biased, with enhanced differentiation potential, while MMP^high^ HSCs are lymphoid-biased with diminished differentiation potential. Mechanistically, low mitochondrial activity upregulates Phosphoinositide 3-kinase (PI3K) signaling to promote HSC differentiation. These insights into the initiation of metabolic heterogeneity could be leveraged to isolate HSCs to efficiently generate desired lineages.

**Highlights:**

- Mitochondrial changes from HSC emergence to maturation govern embryonic hematopoiesis.
- Lowering mitochondrial activity promotes Wnt-dependent HSC expansion and PI3K-dependent differentiation.
- Mitochondrial membrane potential determines lympho-myeloid fate choice in AGM HSCs.
- Aberrant mitochondrial activity in the AGM HSCs perturbs adult hematopoiesis.

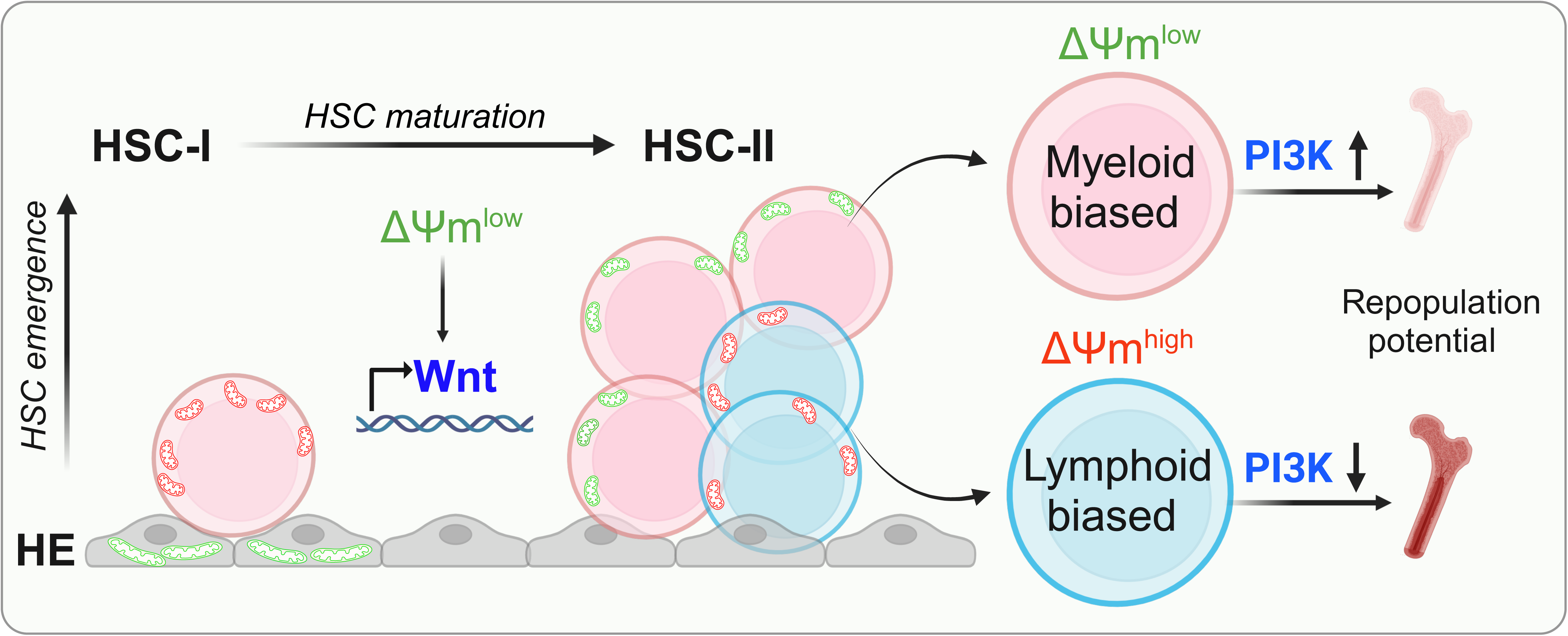

## Introduction

Bone marrow (BM) transplantation relies on the self-renewal and multilineage engraftment capacity of hematopoietic stem cells (HSCs)^1^. However, strategies for expanding or enriching long-term reconstituting HSCs are inefficient, likely due to their inherent functional heterogeneity^2–4^. Therefore, understanding factors governing the ontogeny of HSC heterogeneity is critical for the robust generation of blood progenitors *in-vitro*. Amongst several cell-intrinsic regulators like genetic and epigenetic factors, the mitochondrial metabolic profile reflects cellular dynamics and provides a reliable readout of the cell state^5–7^.

Mitochondrial homeostasis is imperative for the efficient functioning of bone marrow (BM) HSCs^8–10^. Quiescent long-term (LT) HSCs in the BM harbor short, fragmented mitochondria with reduced mitochondrial activity, and generate low reactive oxygen species (ROS). An increase in ROS, oxidative phosphorylation, and mitochondrial biogenesis marks exit from quiescence^9^. Along with its role in HSC fate determination, mitochondrial membrane potential (MMP) and morphology affect BM-HSC fate and give rise to functional heterogeneity in HSCs^11,12^. Pharmacological modulation of mitochondrial activity successfully ameliorates HSC aging-like changes in the murine model^13^.

Several mitochondrial proteins act as scaffolds for signaling complexes and play an important role in maintaining mitochondrial structure and integrity. Altering function of such mitochondria-associated proteins can lead to mitochondrial dysfunction and increases the probability of developing malignant disorders. For example, loss of the mitochondrial fission regulator Dynamin-related protein 1 (Drp1) leads to reduced HSC regenerative potential^14^, while depletion of the fusion protein Mitofusin2 (Mfn2) affects the lymphoid potential^15^. Depletion of the conserved inner mitochondrial membrane protein Asrij/OCIAD1 (Ovarian Carcinoma Immunoreactive Antigen Domain containing protein 1)^16,17^ results in the accumulation of defective mitochondria and increased ROS, leading to loss of HSC quiescence, myeloproliferative and aging-like changes in the HSC pool^18,19^. However, it is unknown when and how this mitochondrial regulation of hematopoiesis originates, and whether it affects early HSC fate and lineage bias.

Definitive HSCs first arise during embryonic days (E) E9.5 to E11.5 in the mouse, through an endothelial to hematopoietic transition (EHT)^20–23^. Activation of a hematopoietic transcriptional program in a subset of endothelial cells of the dorsal aorta in the Aorta-Gonad-Mesonephros (AGM) region, gives rise to a transient precursor called the hemogenic endothelium (HE)^24–27^. The flat endothelial cells change morphology to emerge as rounded cells with a hematopoietic fate, forming the intra-aortic hematopoietic clusters (IAHCs). IAHCs constitute the pre-HSC type-I, which proliferate to form pre-HSC type-II^28^. The latter migrate to the fetal liver, where they undergo expansion around E13.5, and eventually colonize the bone marrow before birth^29,30^. Recent single-cell transcriptomic analyses reveal the upregulation of mitochondrial metabolic pathways upon hematopoietic induction^31^. *In-vitro* human EHT studies have shown that mitochondrial pyruvate and glutamine metabolism are essential for early HSC lineage commitment^32,33^. Due to the rapid flux of metabolites during EHT, subtle differences in mitochondrial properties will likely impact their differentiation. *In-vivo* studies with the murine model have reported a switch from glycolysis to oxidative phosphorylation upon HSC emergence^34,35^. However, a detailed *in-vivo* characterization of mitochondrial parameters during HSC emergence and maturation is lacking. Also, whether and how perturbations in metabolic regulation of the embryonic HSC pool could impact adult hematopoiesis, is not known.

Here, using a multiplexed approach, we show that dynamic changes in mitochondrial activity accompany EHT during the earliest stages of HSC emergence and maturation in the AGM. Using pharmacological and genetic manipulations and HSC transplantation, we ascertained that mitochondrial activity status determines HSC emergence and defines the potency of mature functional HSCs. Further, using bulk and single-cell transcriptomic analyses, we identify key signaling pathways that control HSC maturation and cell fate determination. Thus, our study demonstrates that mitochondrial heterogeneity and lineage choice reported earlier in the bone marrow HSCs originates during the earliest stages of definitive hematopoiesis. Further, using the *asrij* knockout (KO) mouse model, we show that alterations in mitochondrial membrane potential during embryonic stages can manifest as a myeloproliferative disorder in the adult. Our study, therefore, provides a developmental basis for mitochondrial regulation of hematopoietic heterogeneity.

## Results

### Hematopoietic stem cell emergence involves mitochondrial remodeling

To comprehensively understand mitochondrial remodeling during EHT, we assayed the aortic hemogenic endothelium (HE) and emerging HSCs for multiple mitochondrial parameters. *In-situ* visualization of mitochondria in embryonic HSCs was done using R26-Mito-EGFP embryos (a transgenic reporter system expressing mitochondrion-targeted EGFP) and Tom20 as a mitochondrial marker in C57BL/6J wild-type (WT) embryos. The c-kit^+^ emerging AGM HSCs showed higher mitochondrial content than the HE at embryonic day (E) 11.5 (Figure 1A-B). Flow cytometry-based analysis of mitochondrial content was done between HE (CD45^-^ c-kit^-^ CD31^+^) and pre-HSC type-I (CD45^-^ c-kit^+^ CD31^+^) (Figure S1A) using E11.5 R26-mito-EGFP embryos, by comparing the median fluorescence intensity (MFI) of EGFP. Type-I HSCs showed higher mitochondrial content than HE (Figure 1C), and similar results were obtained with E9.5 and E10.5 AGM (Figure S1B-C). Increased mitochondrial content in HSCs was also confirmed by flow cytometry analysis of Tom20 and MitoTracker Deep Red in E11.5 WT AGM (Figure S1D-E). Reverse transcription-quantitative polymerase chain reaction (RT-qPCR) analysis showed that genes involved in mitochondrial biogenesis, such as *Tfam*, *Idh,* and *Pgc1α,* were upregulated in AGM pre-HSC type-I compared to HE (Figure 1D), suggesting that enhanced mitochondrial biogenesis leads to higher mitochondrial mass in the budding HSCs.

**Figure 1.**
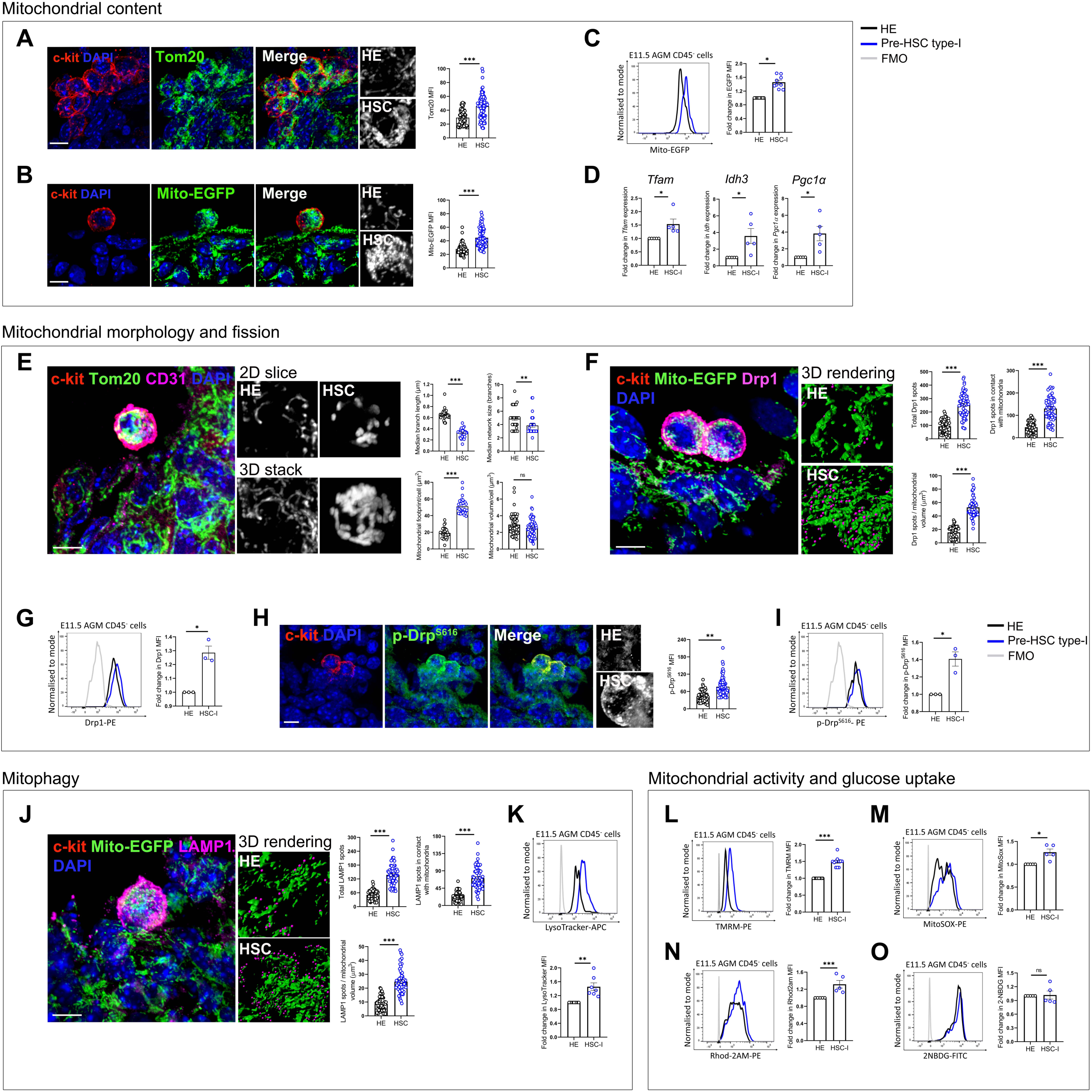
Hematopoietic stem cell emergence involves mitochondrial remodeling. (A-B) Representative immunofluorescence images of mitochondria (green) stained using Tom20 in WT embryos (A) or R26-mito-EGFP expressing embryos (B) in c-kit^+^ HSC clusters (red) in E11.5 AGM sections. Nuclei (blue) were stained using DAPI. Insets show a magnified view of mitochondria (greyscale) in HE and HSC. Graphs show the median fluorescence intensity (MFI) of Tom20 or EGFP, as indicated. Each data point represents an individual cell. Three independent experiments were done (N = 3 embryos); n = 25 HE and 25 HSC for each experiment. Data represent mean ± standard error of mean (SEM). Scale bar = 5µm. (C) Representative flow cytometry histogram shows EGFP intensity in HE (black) and pre-HSC type-I (HSC-I; blue) in E11.5 R26-mito-EGFP AGM. Graph shows fold change in EGFP median fluorescence intensity (MFI) in HSC-I compared to HE. Data are from seven independent experiments, with 7-8 embryos pooled for each experiment (N = 7; n ≥ 7 embryos). Data represent mean ± SEM. (D) RT-qPCR analysis of genes involved in mitochondrial biogenesis-*Tfam*, *Idh*, and *Pgc1α* normalized to the expression of *B2m* in WT E11.5 HE and HSC-I. Fold change in mRNA expression level has been plotted for pre-HSC type-I (HSC-I) with respect to HE. N = 5 independent experiments, mean ± SEM. (E) Representative immunofluorescence image shows Tom20 stained mitochondria (green) in c-kit^+^ HSCs (red) of WT AGM at E11.5, compared to CD31^+^ HE. Magnified insets (greyscale) show a 2D slice view or 3D stack, as indicated. Mitochondrial morphology parameters such as median branch length and network size, footprint, and volume were quantified for HE and HSC. Each data point represents an individual cell. N = 3 embryos, n = 10 HE, and 10 HSC for each experiment. Data represent mean ± SEM. Scale bar = 5µm. (F) Representative immunofluorescence image shows Drp1 (magenta) distribution on the mitochondrial surface (green) within c-kit^+^ HSCs (red) in AGM sections of E11.5 R26-mito-EGFP embryos. Insets show a 3D rendering of Drp1^+^ spots on the mitochondrial surface, done using Imaris software. Total Drp1 spots, Drp1 spots in close contact with a mitochondrial surface (within 0.3µm), and Drp1 spots per unit mitochondrial volume were quantified. Each data point represents an individual cell. N = 3 embryos, n = 20 HE, and 20 HSC for each experiment. Data represent mean ± SEM. (G) Flow cytometry histogram shows Drp1 levels in HE (black) and HSC-I (blue) in E11.5 WT embryos. Fold change in the MFI of Drp1 has been plotted. N = 3; n ≥ 7 embryos. Data represent mean ± SEM. (H) Immunofluorescence images show p-Drp1^Ser616^ (green) in c-kit^+^ HSCs (red) of E11.5 WT AGM sections. Magnified insets show p-Drp1^Ser616^ intensity (greyscale) in HE and HSC. Fold change in p-Drp1^Ser616^ intensity has been plotted for HSC-I with respect to HE. Each data point represents an individual cell. N = 3 embryos; n = 25 HE and 25 HSC for each experiment. Data represent mean ± SEM. Scale bar is 5µm. (I) Flow cytometry histogram shows p-Drp1^S616^ levels in HE (black) and HSC-I (blue) of E11.5 WT embryos. Fold change in MFI of p-Drp1 has been plotted. N = 3; n ≥ 7 embryos, and data represent mean ± SEM. (J) Immunofluorescence images show the distribution of LAMP1^+^ lysosomes (magenta) on the mitochondrial surface (green) within c-kit^+^ HSCs (red) in E11.5 R26-mito-EGFP AGM sections. Insets show a 3D rendering of LAMP1^+^ spots on the mitochondrial surface, done using Imaris software. Total LAMP1 spots, LAMP1 spots in close contact with a mitochondrial surface (within 0.3 µm), and LAMP1 spots per unit mitochondrial volume were quantified. N = 3 embryos, n = 20 HE, and 20 HSC for each experiment. Data represent mean ± SEM. (K) Flow cytometry histogram shows LysoTracker-Deep Red intensity in HE (black) and HSC-I (blue) of E11.5 WT embryos. Graph shows fold change in LysoTracker MFI. N = 7, n ≥ 7 embryos, and data represent mean ± SEM. (L-N) Flow cytometry histograms show mitochondrial activity parameters. Mitochondrial membrane potential (MMP) was measured using TMRM (L), reactive oxygen species (ROS) was measured using MitoSOX Red (M), and mitochondrial calcium was measured using Rhod-2 AM (N) in E11.5 WT AGM. Graphs show fold change in MFI of mitochondrial probes, N ≥ 5, n ≥ 7 embryos, and data represent mean ± SEM. (O) Flow cytometry histogram shows the intensity of 2-NBDG in HE (black) and HSC-I (blue) of E11.5 WT embryos. Graph shows fold change in 2-NBDG MFI. N = 5, n ≥ 7, and data represent mean ± SEM.

Mitochondrial morphology informs about the cellular metabolic state, and mitochondrial fission-fusion dynamics are known to affect HSC or progenitor potency and lineage choice^14,36^. Morphometric analysis of the mitochondrial network was performed in Tom20 stained E11.5 WT AGM sections in c-kit^+^ HSCs using high-resolution confocal microscopy. There was no change in the overall mitochondrial volume per cell between HE and HSC. However, while HE had long tubular mitochondria, with increased branch length and network size, HSCs harbored short, fragmented mitochondria, indicating mitochondrial fission (Figure 1E). Mitochondrial fission is essential for the maintenance of the mitochondrial network and relies on Dynamin-related protein 1 (Drp1). Upon activation, Drp1 gets phosphorylated and is recruited to the mitochondrial surface, where it carries out fission via its GTPase activity^37^. Three-dimensional (3D) rendering of Drp1 to analyze contact with the mitochondrial surface using R26-mito-EGFP AGM sections revealed increased localization of Drp1 to the mitochondrial surface in HSCs compared to the HE (Figure 1F). Imaging and flow cytometric analysis showed a significant increase in Drp1 levels (Figure 1G) in budding HSCs, with a corresponding increase in its activated form, phosphorylated-Drp1^S616^ (Figure 1H-I), confirming higher mitochondrial fission in emerging HSCs. A similar analysis was done with the lysosomal marker Lysosomal-associated membrane protein 1 (LAMP1) to assess for mitophagy, which is known to increase in cells with high mitochondrial fragmentation. LAMP1^+^ lysosomes showed increased association with the mitochondrial surface in HSCs compared to HE (Figure 1J), along with an increase in total lysosomal content (Figure 1K), indicating higher mitophagy, which correlates with the high rate of fission in HSCs.

Changes in mitochondrial mass and morphology could indicate changes in mitochondrial activity. Live analysis of mitochondrial metabolic status was done using fluorescent mitochondrial activity probes. Flow cytometry-based analysis of tetramethylrhodamine methyl ester (TMRM), that indicates mitochondrial membrane potential (MMP) showed higher levels in pre-HSC type-I compared to HE at E11.5 (Figure 1L) as well as during EHT initiation at E9.5 and E10.5 (Figure S1F-G). Mitochondrial reactive oxygen species (ROS) assayed using MitoSOX Red and calcium signaling levels assayed using Rhod-2 AM were higher in Pre HSC-type-I compared to the HE (Figure 1M-N). Higher MMP, ROS, and calcium indicate an overall increase in mitochondrial activity in emerging HSCs. Assaying for glycolytic capacity using 2-(N-(7-Nitrobenz-2-oxa-1,3-diazol-4-yl)Amino)-2-Deoxyglucose (2-NBDG) showed no change in glucose uptake between HE and pre-HSC type-I (Figure 1O), indicating that not glycolysis, but mitochondrial respiration could be the predominant source of energy production during the early phase of EHT.

### HSC maturation is accompanied by an increase in mitochondrial activity and MMP-based segregation of mature HSCs

Emergent pre-HSC type-I undergo maturation to form definitive embryonic HSCs. However, changes in mitochondrial dynamics and metabolic state accompanying the transition of nascent embryonic HSCs to mature functional HSCs *in-vivo* are unknown. Metabolic readouts were assayed between pre-HSC type-I and pre-HSC type-II (CD31^+^ c-kit^+^ CD45^+^) at E11.5 to analyze the changes in mitochondrial activity during HSC maturation. Flow cytometric analysis using R26-Mito-eGFP embryos revealed a decrease in mitochondrial mass upon transition to pre-HSC type-II (Figure 2A). RT-qPCR analysis further confirmed the downregulation of genes involved in mitochondrial biogenesis like *Tfam, Idh, and Pgc1a* (Figure 2B). Lysosomal content was also increased in type-II HSCs, which could indicate higher mitophagy (Figure 2C). Along with mitochondrial mass, mitochondrial ROS (Figure 2D) and calcium levels (Figure 2E) were also reduced in type-II HSCs compared to type-I HSCs.

**Figure 2.**
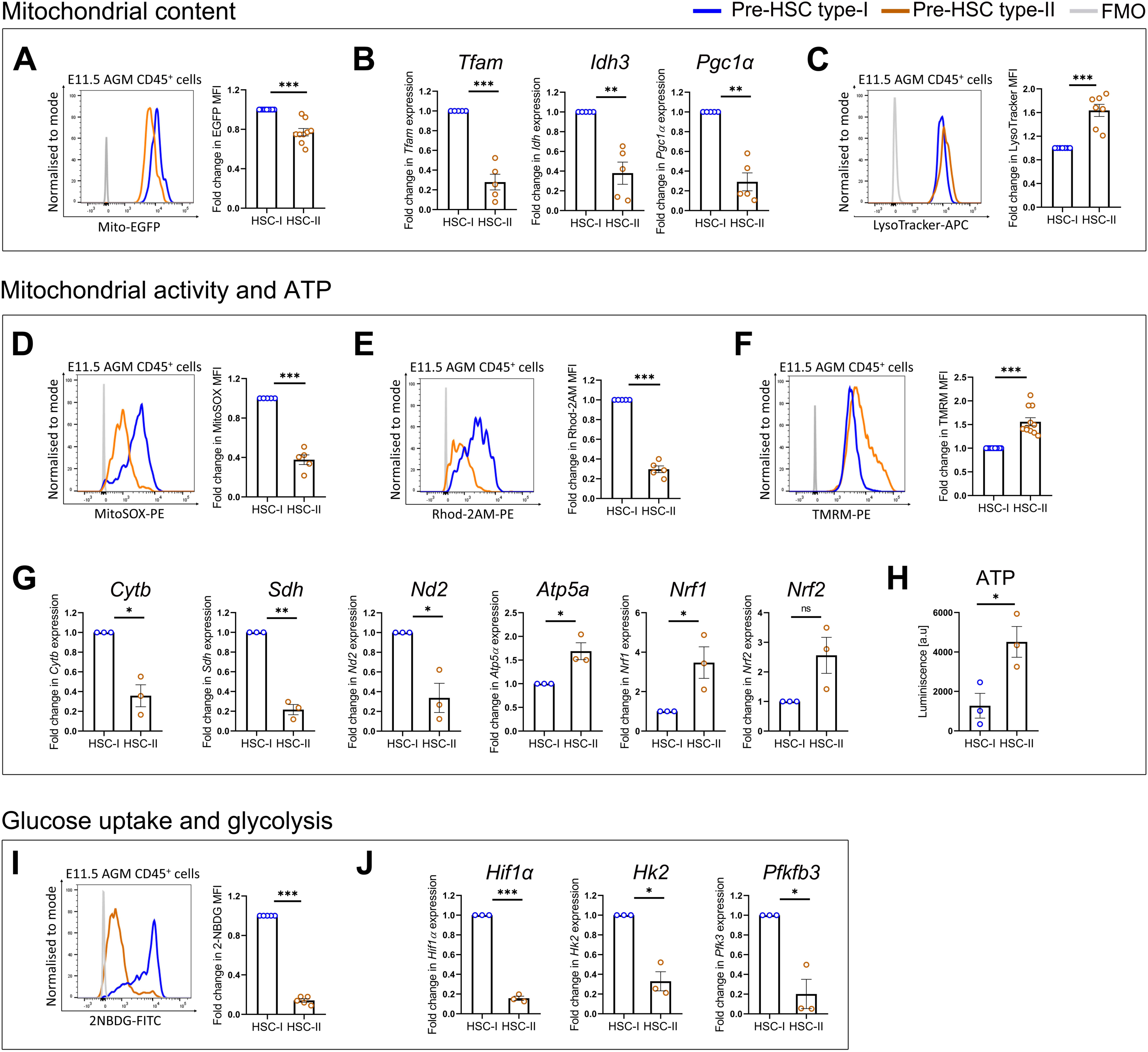
HSC maturation is accompanied by an increase in mitochondrial activity, with MMP-based segregation of mature HSCs. (A) Representative flow cytometry histogram shows EGFP intensity in pre-HSC type-I (HSC-I; blue) and pre-HSC type-II (HSC-II; orange) of E11.5 R26-mito-EGFP embryos. Graph shows fold change in EGFP MFI. N = 9, n ≥ 7 embryos, and data represent mean ± SEM. (B) RT-qPCR analysis of genes involved in mitochondrial biogenesis-*Tfam*, *Idh*, and *Pgc1α*, normalized to the expression of *B2m* in WT E11.5 HSC-I and HSC-II. Fold change in mRNA expression level has been plotted for HSC-II with respect to HSC-I. N = 5 independent experiments, data represent mean ± SEM. (C) Flow cytometry histogram shows Lysotracker-Deep Red intensity in HSC-I and HSC-II of E11.5 WT embryos. Graph shows fold change in LysoTracker MFI. N = 7, n ≥ 7 embryos, and data represent mean ± SEM. (D-F) Representative flow cytometry histograms for MitoSOX (D), Rhod-2 AM (E), and TMRM (F) intensity in HSC-I and HSC-II of E11.5 WT embryos. Graphs show fold change in MFI. N ≥ 5, n ≥ 7 embryos, and data represent mean ± SEM. (G) RT-qPCR analysis of genes involved in mitochondrial respiration-*Cytb, Sdh, Nd2, Atp5α, Nrf1, and Nrf2,* normalized to *B2m* expression in WT E11.5 HSC-I and HSC-II. Fold change in mRNA expression level has been plotted for HSC-II with respect to HSC-I. N = 3 independent experiments, data represent mean ± SEM. (H) Graph shows ATP content (luminescence) in sorted HSC-I and HSC-II cells from E11.5 WT embryos. N = 3, n ≥ 7 embryos, data represent mean ± SEM. (I) Flow cytometry histogram shows the intensity of 2-NBDG in HSC-I and HSC-II of E11.5 WT embryos. Graph shows fold change in 2-NBDG MFI. N = 5, n ≥ 7, and data represent mean ± SEM. (J) RT-qPCR analysis of genes involved in glycolysis-*Hif1α, Hk2, and Pfk3,* normalized to *B2m* expression, in WT E11.5 HSC-I and HSC-II. Graph shows fold change in mRNA expression level in HSC-II with respect to HSC-I. N = 3 independent experiments, data represent mean ± SEM.

In contrast to the reduced mitochondrial content, MMP was increased upon transition to type-II HSCs (Figure 2F). Interestingly, RT-qPCR analysis of genes involved in mitochondrial respiration showed downregulation of *Cox1, Sdh, Nd2, Cytb,* and upregulation of *Atp5a*, *Nrf1,* and *Nrf2* in type-II HSCs compared to type-I (Figure 2G). Analysis of AGM single-cell transcriptome data^38^ also revealed heterogeneity in the expression of key mitochondrial genes within a given subset of mature embryonic HSCs (Figure S2A). Also, the histogram for TMRM intensity profile showed a small population with high MMP, which separated from the bulk of the population, indicating that the type-II HSC pool is heterogeneous in terms of MMP (Figure S2B). Similar results were obtained at E10.5 AGM (Figure S2C), suggesting a conserved pattern of MMP heterogeneity in pre-HSC type-II.

Cellular ATP levels were assayed to test whether an overall increase in mitochondrial activity leads to higher energy production. Type-II HSCs showed a drastic increase in total ATP levels (Figure 2H), compared to type-I. 2-NBDG-based analysis of glucose uptake revealed reduced glycolytic capacity in the type-II HSCs compared to type-I (Figure 2I), with a concomitant decrease in expression levels of genes involved in glycolysis such as *Hk2*, *Pfk2* and *Hif1α* (Figure 2J). Downregulation of glycolysis in mature HSCs further confirms that mitochondrial respiration is the predominant source of energy production at this stage. Even though a majority of the type-II HSCs have low mitochondrial activity, the subset of cells with high MMP seem sufficient to effect an overall increase in energy metabolism.

### Mitochondrial activity controls HSC maturation, but not emergence

We showed that mitochondrial activity gradually increases during EHT, and mature HSCs differ in their level of mitochondrial activity. Therefore, *ex-vivo* AGM explant cultures were used to test the impact of mitochondrial modulation on EHT. E10.5 and E11.5 WT AGM explants were cultured for 48 hours in the presence of carbonyl cyanide m-chlorophenylhydrazine (CCCP), which decreases mitochondrial activity by electron transport chain (ETC) uncoupling, or Mitoquinol (MitoQ) which quenches ROS, resulting in increased mitochondrial activity (Figure 3A). After 48 hours of treatment, frequencies of HE, pre-HSC type-I, and type-II were analyzed to identify the cell type most impacted by mitochondrial modulation. There was no change in HE and pre-HSC type-I, in the presence of CCCP and MitoQ for E10.5 and E11.5 AGM. However, pre-HSC type-II increased drastically in frequency upon CCCP treatment for both stages (Figure 3B). Interestingly, type-II HSC frequency from the E10.5 explant was unchanged in the presence of MitoQ, while for E11.5, there was a substantial decrease in pre-HSC type-II (Figure 3C; Figure S3A-B).

**Figure 3.**
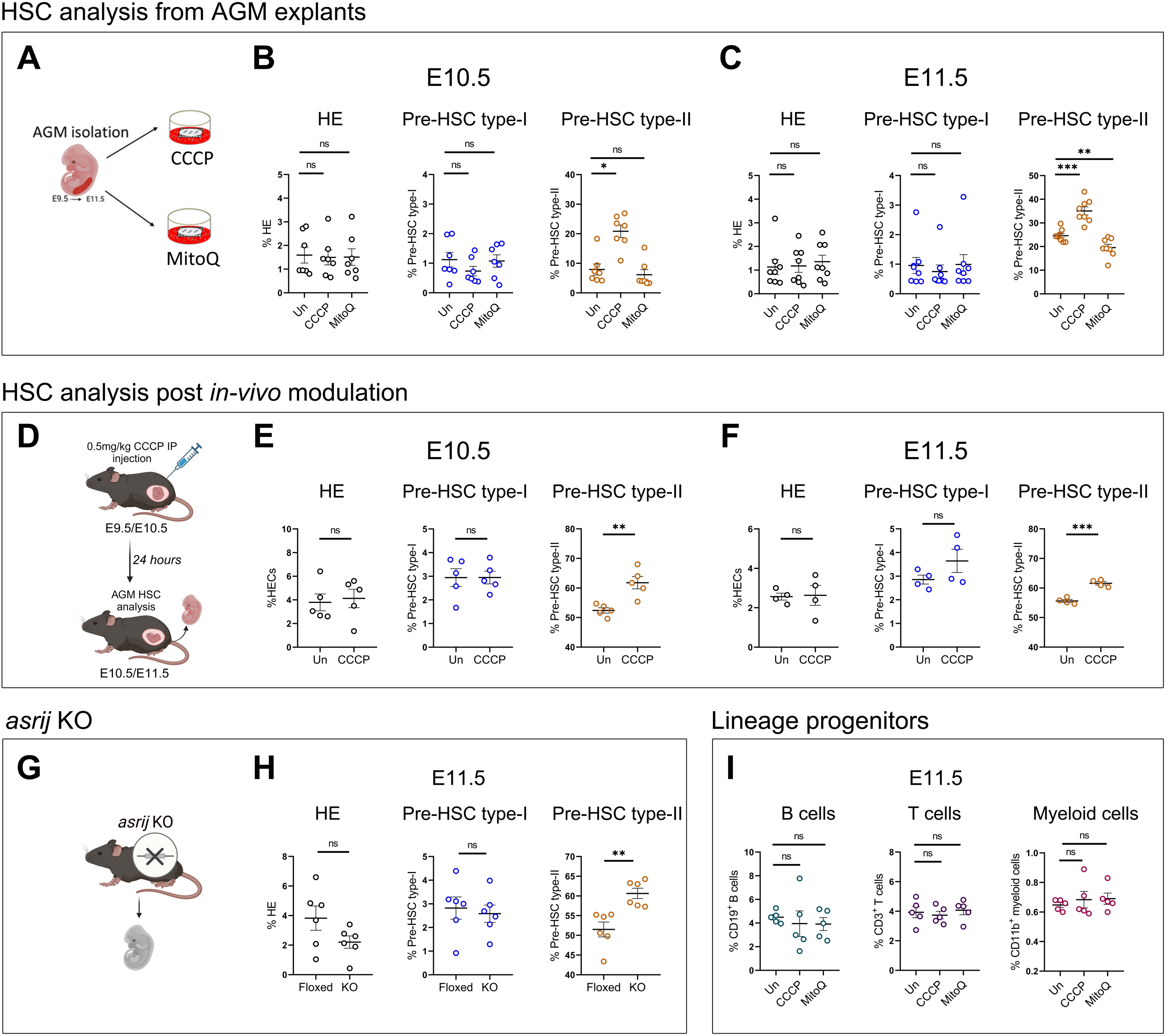
Mitochondrial activity causally regulates HSC maturation, but not specification. (A) Schematic of AGM explant culture experiment with CCCP and Mitoquinol. (B-C) Flow cytometric analysis of HE, pre-HSC type-I, and pre-HSC type-II in WT AGM explants post 48-hour treatment with CCCP or mitoQ at E10.5 (B) and E11.5 (C). N ≥ 7 independent experiments, with n = 3 AGM per experiment/condition. Un=untreated, data represent mean ± SEM. (D) Schematic of *in-vivo* CCCP treatment of WT pregnant dams at E9.5 or E10.5 for analysis post 24 hours. (E-F) Flow cytometric analysis of HE, pre-HSC type-I, and pre-HSC type-II from CCCP treated or control (Un) WT mice at E10.5 (E) and E11.5 (F). N = 4, with n ≥ 7 embryos per experiment, data represent mean ± SEM. (G) Representative schematic of *asrij* KO mouse. (H) Flow cytometric analysis of HE, pre-HSC type-I, and pre-HSC type-II from E11.5 control (floxed) and *asrij* KO embryos (F). N = 6, n ≥ 7 embryos per experiment, data represent mean ± SEM. (I) Flow cytometric analysis of B cell, T cell, and myeloid cell lineages in WT E11.5 AGM explants post 72-hour treatment with CCCP or mitoQ. N = 5, n = 3 AGM per experiment/condition. Un = untreated, data represent mean ± SEM.

To test whether mitochondrial modulation has a similar effect *in-vivo*, pregnant dams were treated with CCCP by intraperitoneal (IP) injection 24 hours before assaying for HSC frequencies at E10.5 and E11.5 (Figure 3D). As expected, CCCP treated embryos showed an expanded pre-HSC type-II pool with no effect on HE and pre-HSC type-I at E10.5 and E11.5 (Figure 3E-F; Figure S3C).

In addition to the pharmacological modulation of mitochondrial activity, we used the *asrij* KO mouse as a genetic model of perturbed mitochondrial homeostasis (Figure 3G). Flow cytometry analysis revealed lower MMP in *asrij* KO AGM HSCs (Figure S3D) compared to the control. Since this model exhibits lower mitochondrial activity, we hypothesized an effect similar to CCCP treatment. As expected, KO AGM at E11.5 showed expansion of the pre-HSC type-II pool with no change in the frequencies of HE and type-I HSCs (Figure 3H; Figure S3E).

We further checked the effect of mitochondrial modulation on lineage-committed progenitors. E11.5 AGM explants were cultured for 72 hours in the presence of CCCP and MitoQ to analyze the lymphoid and myeloid progenitor frequencies. CD11b was used as a marker for early myeloid lineage cells, while CD19 and CD3 were used for B and T cell lymphoid lineages, respectively. Surprisingly, there was no significant change in the frequencies of lineage committed progenitors in the presence of CCCP and MitoQ compared to the untreated controls (Figure 3I; S3H). Thus, a change in mitochondrial activity affects HSC maturation but does not affect hematopoietic specification and downstream lineage commitment.

Decreasing mitochondrial activity led to enhanced production of type-II HSCs at both E10.5 and E11.5. Increasing the mitochondrial activity, however, only impacts E11.5 type-II HSCs. This shows that mitochondrial homeostasis fine-tunes embryonic HSC development. Further, mitochondrial activity inversely regulates mature HSC production, exerting a stage-specific and temporally regulated control on EHT.

### Lower mitochondrial activity upregulates Wnt signaling to promote HSC maturation

To gain mechanistic insights into the mitochondrial regulation of embryonic HSC maturation, bulk RNA sequencing was performed in WT E10.5 control and CCCP-treated HSCs. Magnetic-activated cell sorting (MACS) was used to isolate c-kit^+^ HSCs from AGM explants post 48 hours of treatment with CCCP. Principal component analysis (PCA) of untreated and CCCP-treated samples showed 65% transcriptional separation, with a 22% separation observed across replicates of the same sample (Figure S4A). Analysis of differentially expressed genes (DEGs) using the DESeq2 package in the R program revealed 1201 DEGs (p-value < 0.05, log2fold change > 0.5 or < -0.5) in CCCP-treated HSCs compared to control, with 1056 genes downregulated and 145 genes upregulated (Figure 4A; Figure S4B; Table S2). The heatmap of all significant DEGs revealed the presence of distinct transcriptional subsets across control and CCCP-treated HSCs (Figure 4B). Gene Ontology (GO) analysis was done to identify the biological processes that are differentially regulated. Cell fate commitment, translation, post-transcriptional gene regulation, and biosynthetic processes were most significantly upregulated (Figure 4C) in the CCCP-treated HSCs, while most metabolic processes were downregulated (Figure 4D). Expression of genes involved in cell proliferation was unchanged (Figure S4C). Gene ontology (GO) revealed an enrichment of the Wnt signaling pathway, gonadotropin-releasing hormone receptor signaling pathway, and cadherin signaling pathway (Figure 4E), with the highest enrichment of Wnt signaling (Figure 4F). This is accompanied by a concomitant decrease in the expression of genes involved in the TGF-β pathway (Figure S4D-E), whose upregulation is known to inhibit EHT^39^.

**Figure 4.**
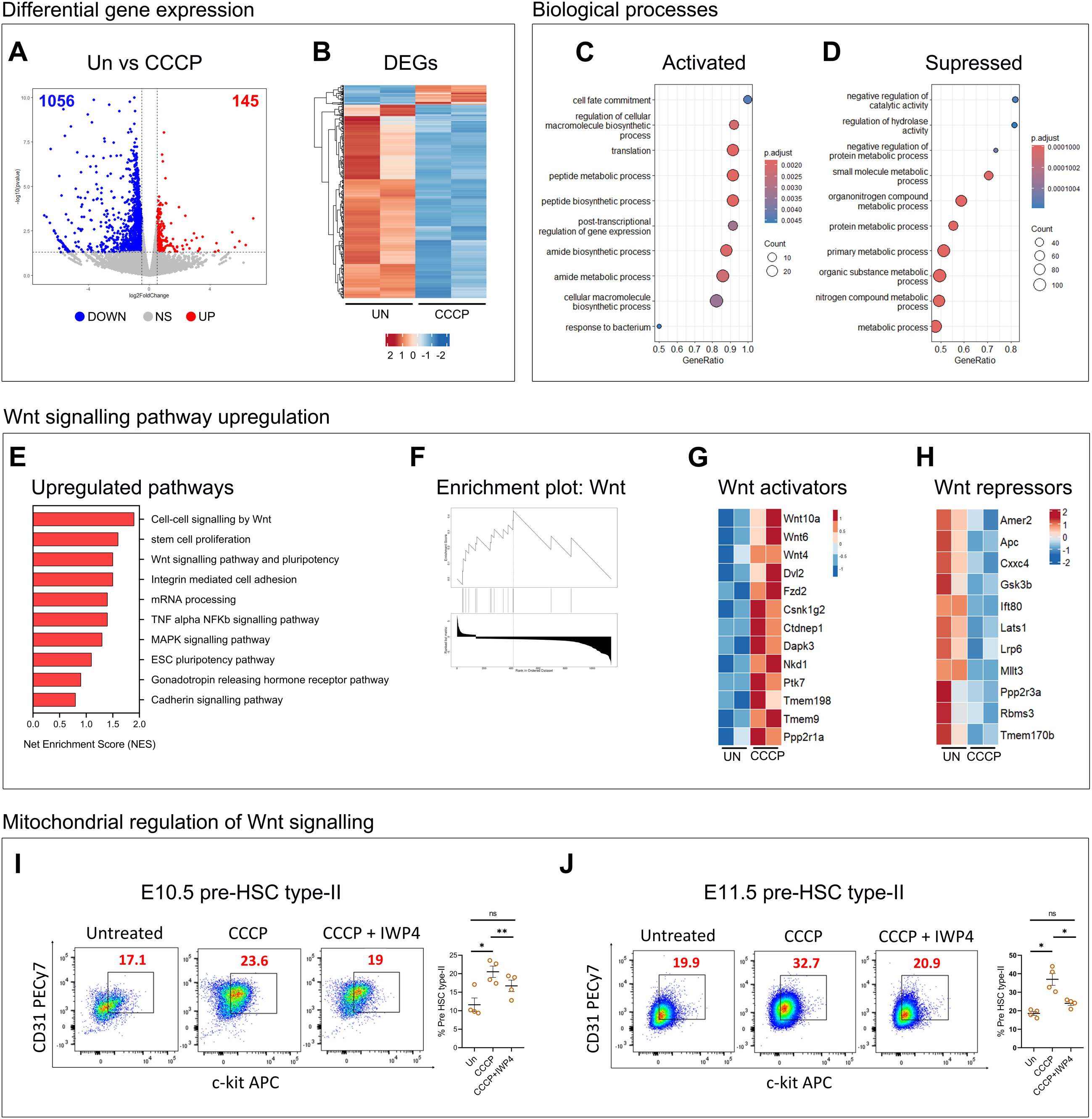
Lower mitochondrial activity upregulates Wnt signaling to promote HSC maturation. (A) Volcano plot shows differentially expressed genes (DEGs) in E10.5 AGM explant CCCP treated HSCs compared to untreated (Un) controls (red = upregulated, blue = downregulated, grey = non-significant; fold change > 0.5, p-value < 0.05) N = 2 independent experiments. See Table S2 for the gene list. (B) Heatmap represents all significant DEGs in E10.5 AGM explant CCCP treated HSCs compared to untreated (Un) controls. (C-D) Dot plots represent biological processes significantly activated (C) or repressed (D) in E10.5 AGM explant CCCP-treated HSCs compared to untreated controls. The dot size represents the gene counts for each biological process. (E) Plot shows net enrichment score (NES) for signaling pathways upregulated in E10.5 AGM explant CCCP treated HSCs compared to untreated controls. (F) GSEA enrichment plot for Wnt signaling shows enrichment score or rank of Wnt signaling across the ordered dataset in E10.5 AGM explant CCCP treated HSCs compared to untreated controls. (G-H) Heatmap shows expression of genes involved in Wnt pathway activation (G) or repression (H) between control and CCCP-treated E10.5 AGM explant HSCs. See Table S3 for normalized gene counts. (I-J) Flow cytometry analysis of HE, pre-HSC type-I, and pre-HSC type-II in AGM explants post 48-hour treatment with CCCP or CCCP + IWP at E10.5 (B) and E11.5 (C). N = 4 independent experiments, with n = 3 AGM per experiment/condition. Un=untreated, data represent mean ± SEM.

We chose to analyze the role of the Wnt signaling pathway, as it promotes definitive hematopoiesis and actively regulates HSC function in the embryo^40^. Genes involved in the canonical Wnt signaling pathway, like the ligands *Wnt4*, *Wnt6*, and *Wnt10a* and receptors *Frizzled* and *Dishevelled*, are upregulated in CCCP-treated HSCs. Further, Wnt pathway inhibitors such as *Apc* and *GSK3*β, which are components of the β-catenin destruction complex, are downregulated (Figure 4G-H; Table S3).

To confirm that mitochondrial activity promotes expansion of the mature HSC pool via the Wnt signaling axis, E10.5, and E11.5 AGM explants were treated with Inhibitor of Wnt production-4 (IWP-4), a Wnt signaling antagonist, in the presence of CCCP. While the CCCP-treated samples had an expanded pre-HSC type-II pool (Figure 4I-J), the type-II HSC pool in IWP + CCCP treated AGM was comparable to the untreated AGM, while HE and pre-HSC type-I remained unaffected at E10.5 and E11.5 (Figure S4E-F). Thus, Wnt pathway inhibition under conditions of reduced mitochondrial activity restricts type-II HSC expansion. Therefore, during HSC maturation, lowering the mitochondrial activity in the AGM drives the production of definitive HSCs by activating the Wnt signaling cascade. Our results establish a novel axis of metabolic regulation of embryonic HSC development through the canonical Wnt signaling pathway.

### Mitochondrial membrane potential determines pre-HSC type-II fate

Mitochondrial membrane potential (MMP) is a well-established determinant of functional heterogeneity in adult BM LT-HSCs^13,41,42^. The fluorescence intensity profile of TMRM in pre-HSC type-II (Figure S2B-C) suggests an underlying MMP-based heterogeneity in definitive embryonic HSCs. Analysis of MMP in the WT AGM hematopoietic pool showed that HE and type-I HSCs did not exhibit any marked segregation based on TMRM intensity. The type-II HSCs, however, could be segregated into a major pool of low MMP (MMP^low^) and a smaller fraction (around 15%) with high MMP (MMP^high^), starting from E10.5 (Figure S5A) and more distinctly at E11.5 (Figure 5A-B). Though MMP is highly elevated in the MMP^high^ subset (Figure 5C), the mitochondrial mass was unchanged between the MMP^low^ and MMP^high^ cells (Figure 5D). We next asked whether this distribution leads to mitochondrial activity-driven functional differences in pre-HSC type-II.

**Figure 5.**
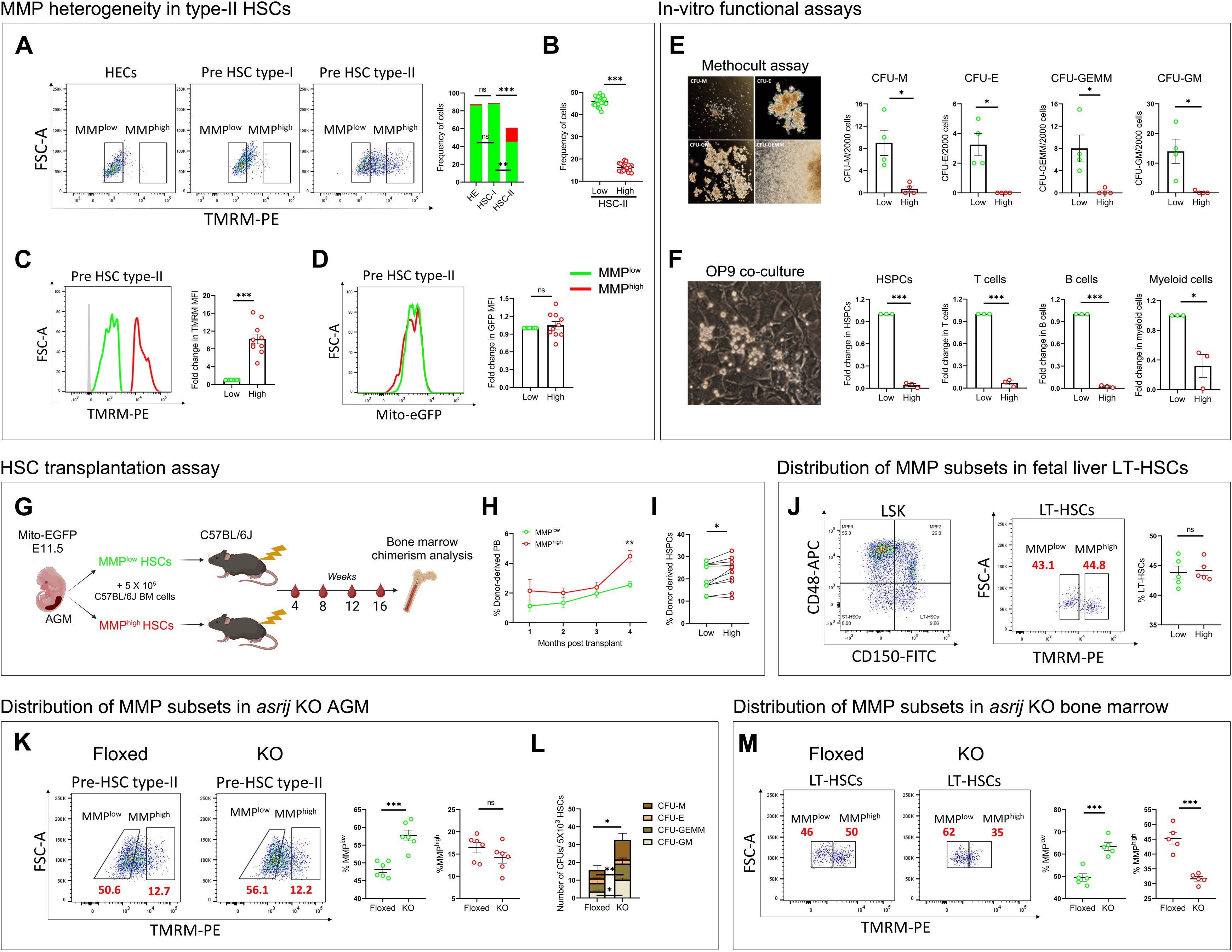
Mitochondrial membrane potential determines pre-HSC type-II fate. (A) Representative flow cytometry scatter plots show TMRM-based cellular distribution in WT E11.5 HE, pre-HSC type-I, and pre-HSC type-II. Graph shows frequency of cells with low and high mitochondrial activity in each cell type. N = 20, n ≥ 7 embryos, data represent mean ± SEM. (B) Graph shows the frequency of MMP^low^ (Low) and MMP^high^ (High) cells in WT E11.5 pre-HSC type-II. N = 20, n ≥ 7 embryos, data represent mean ± SEM. (C) Flow cytometry histogram shows TMRM intensity in WT E11.5 HSC-II MMP^low^ (Low) and MMP^high^ (High) subsets. Graph shows fold change in TMRM MFI. N = 10, n ≥ 7 embryos, data represent mean ± SEM. (D) Flow cytometry histogram shows EGFP intensity in MMP^low^ (Low) and MMP^high^ (High) subsets of E11.5 R26-mito-EGFP AGM type-II HSCs. The graph shows fold change in EGFP MFI. N = 10, n ≥ 7 embryos, data represent mean ± SEM. (E) Representative images of *in-vitro* methylcellulose colony forming assay colonies. Graphs show quantification of macrophage colony forming unit (CFU-M), erythroid (CFU-E), granulocyte-macrophage (CFU-GM), and granulocyte erythrocyte monocyte-macrophage (CFU-GEMM) colonies from MMP^low^ (Low) and MMP^high^ (High) subsets post 2 weeks. N = 4, n = 2000 HSCs from ≥ 7 pooled AGM, data represent mean ± SEM. (F) Representative image of OP9 co-culture assay. Graphs show fold change in numbers of HSPCs, T cells, B cells, and myeloid cells post 2 weeks of co-culture in MMP^low^ (Low) and MMP^high^ (High) subsets. N = 3, n = 1000 HSCs from ≥ 7 pooled AGM, data represent mean ± SEM. (G) Schematic representing the HSC transplantation assay workflow. (H) The graph shows the percentage chimerism in peripheral blood of WT irradiated recipients injected with E11.5 R26-mito-EGFP MMP^low^ (Low) or MMP^high^ (High) HSCs at 1, 2, 3, and 4 months post-transplant. N = 11 recipients/condition, data represent mean ± SEM. (I) Graph shows percentage donor chimerism in HSPCs of C57WT irradiated recipients injected with E11.5 R26-mito-EGFP MMP^low^ or MMP^high^ HSCs at 4 months post-transplant. N = 11 recipients/condition, data represent mean ± SEM. (J) Flow cytometry-based gating of WT E15.5 fetal liver Lin^-^ Sca1^+^ c-kit^+^ HSPCs (LSK). The scatter plot and graph show the frequency of MMP subsets in FL LT-HSCs. N = 5, n ≥ 5 fetal livers, data represent mean ± SEM. (K) Representative flow cytometry scatter plots show the distribution of MMP subsets in E11.5 *asrij* floxed and KO pre-HSC type-II. The graph shows the frequency of MMP^low^ and MMP^high^ subsets. N = 5, n ≥ 7 embryos, data represent mean ± SEM. (L) Graph shows quantification of methylcellulose CFUs in *asrij* floxed and KO pre-HSC type-II. N = 4, n = 5000 HSCs from ≥ 7 pooled AGM, data represent mean ± SEM. (M) Representative flow cytometry scatter plots show the distribution of MMP subsets in *asrij* floxed and KO bone marrow (4 months). Graph shows frequency of MMP^low^ and MMP^high^ subsets. N = 5, data represent mean ± SEM.

To understand whether MMP determines cell fate decisions, the MMP^low^ and MMP^high^ subsets were sorted in equal numbers using fluorescence-activated cell sorting (FACS), and subjected to *in-vitro* hematopoietic differentiation assays and *in-vivo* HSC transplantation assay. Methylcellulose colony forming assay and OP9 co-culture assay revealed that the MMP^low^ subset has relatively higher *in-vitro* multilineage differentiation potential. However, the MMP^high^ pool was refractile to differentiation, which could indicate higher quiescence (Figure 5E-F). To test the impact of the two MMP pools *in-vivo*, we transplanted ∼500 sorted pre-HSC type-II MMP^low^ and MMP^high^ cells from R26-mito-EGFP embryos into lethally irradiated C57BL/6J wild-type (WT) mice along with WT whole BM support cells (Figure 5G). Peripheral blood and bone marrow HSPC chimerism analysis after 4 months of HSC transplant revealed a higher degree of chimerism in the MMP^high^ subset compared to the MMP^low^ HSCs (Figure 5H-I; Figure S5B). However, the degree of chimerism in other lineage-committed cells was unchanged (Figure S5C). This indicates that mitochondrial activity dictates early HSC fate, where lower MMP promotes higher differentiation, whereas high MMP maintains quiescence, contributing to enhanced long-term HSPC reconstitution.

To further track these subsets during hematopoietic development, long-term HSCs (LT-HSCs) from the WT E15.5 fetal liver were analyzed. Flow cytometry analysis showed that the LT-HSC pool in the fetal liver has an equal distribution of the MMP^low^ and MMP^high^ subsets (Figure 5J). The short-term (ST)-HSC pool, on the other hand, was skewed towards MMP^high^ (Figure S5D), as reported earlier in the bone marrow HSPCs^42^. Analysis of mitochondrial content and potential in the fetal liver HSPC pool also revealed the lowest mitochondrial content and activity in LT-HSCs, with a modest increase along the differentiation hierarchy (Figure S5E-F), much like the bone marrow.

To further validate whether mitochondrial activity levels determine the differentiation potential, we used the *asrij* KO AGM as a genetic model for altered mitochondrial activity. Indeed, flow cytometry analysis revealed an expansion of the MMP^low^ subset in the KO AGM (Figure 5K). Further, there was enhanced *in-vitro* differentiation potential of the pre-HSC type-II pool in KO compared to control, thus confirming that lower MMP promotes differentiation (Figure 5L).

While the distribution of MMP subsets changes through development, it is unclear whether the embryonic MMP status impacts post-natal hematopoiesis. To interrogate this, we examined the *asrij* KO bone marrow LT-HSCs. The embryonic KO HSC pool is skewed to MMP^low^. Bone marrow LT-HSCs of *asrij* KO also showed expansion of the MMP^low^ pool (Figure 5M), leading to loss of BM-HSC quiescence as reported previously^18^. We therefore show that metabolic HSC heterogeneity reported in the bone marrow originates during embryonic hematopoiesis in pre-HSC type-II, attains a BM-like state in the fetal liver, and impacts post-natal hematopoietic homeostasis.

### Mitochondrial activity drives lympho-myeloid lineage bias in embryonic HSCs

To understand the transcriptional basis of MMP-based heterogeneity in the AGM hematopoietic pool, single-cell RNA sequencing (scRNA seq) was performed on MMP^low^ and MMP^high^ FACS sorted cells from E11.5 WT AGM (Figure 6A). Unsupervised hierarchical clustering using known markers of the AGM cell populations (Figure S6A) identified 11 distinct clusters in the MMP^low^ pool, and 8 clusters in the MMP^high^ pool (Figure 6B). Uniform manifold approximation and projection (UMAP) visualization showed that the MMP^low^ population comprised all cell types of the EHT continuum, including vascular endothelium, hemogenic endothelium, EHT cells, intra-aortic hematopoietic cluster (IAHC), pro-HSCs, pre HSC-type-I and pre HSC-type-II. On the other hand, the MMP^high^ pool consisted mainly of the hematopoietic population, including EHT cells, IAHC, pro-HSCs, and Pre HSC-type-I and type-II (Figure 6C; Figure S6B). Comparative marker gene expression analysis of endothelial, EHT, and hematopoietic cells revealed upregulation of hematopoietic genes in the MMP^high^ pool (Figure 6D), indicating that AGM cells with high mitochondrial activity are enriched for hematopoietic cells.

**Figure 6.**
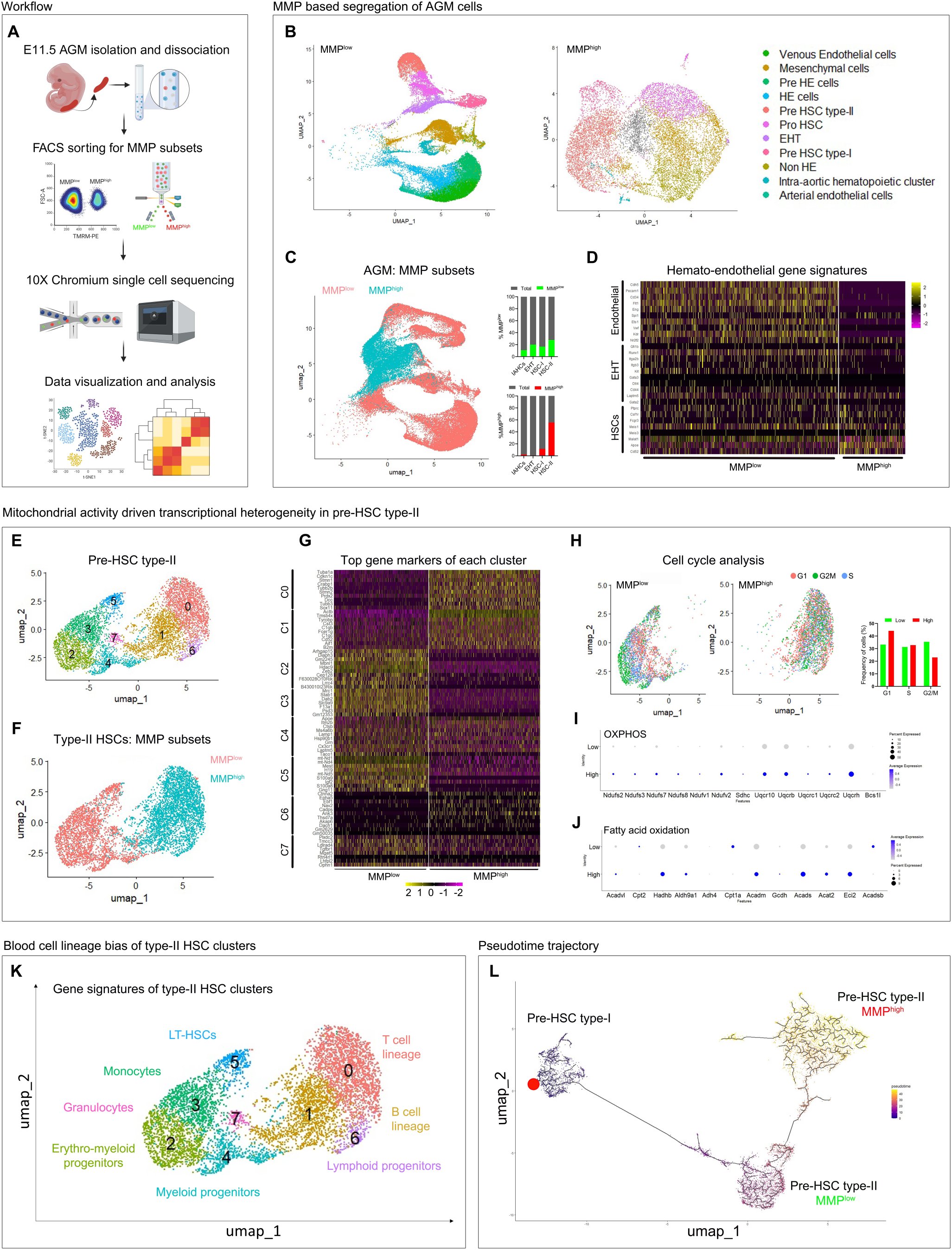
Mitochondrial activity drives lympho-myeloid lineage bias in embryonic HSCs. (A) Schematic representing workflow of single-cell RNA sequencing (scRNA-seq) with FACS sorted MMP^low^ and MMP^high^ cells from WT E11.5 AGM using Chromium 10X single-cell sequencing protocol. (B) Plots show uniform manifold approximation and projection (UMAP) based clustering of MMP^low^ and MMP^high^ cells from WT E11.5 AGM. Analyses represent pooled datasets from two independent experiments. (C) Plot shows UMAP-based clustering of pooled AGM transcriptome, with MMP subsets color-coded. Graph shows distribution of AGM hemato-endothelial cells in the MMP subsets of WT E11.5 AGM. (D) Heatmap representing expression levels of endothelial, EHT, and hematopoietic genes in WT E11.5 AGM MMP subsets. (E) Plot shows UMAP-based clustering of WT E11.5 pre-HSC type-II HSCs from the whole AGM. (F) Plot shows MMP subsets (color-coded) of the pre-HSC type-II clusters. (G) Heatmap shows expression levels of the top marker genes of a cluster in MMP^low^ and MMP^high^ type-II HSC subsets. See Table S4. (H) Plot shows UMAP cell cycle analysis on MMP subsets of WT E11.5 pre-HSC type-II. G1, G2/M, and S phase cells are color-coded, and the graph shows the number of cells in each phase of the cell cycle in the MMP^low^ and MMP^high^ subsets. (I-J) Dot plots show the expression of genes involved in OXPHOS (I) and fatty acid oxidation (FAO; J) in MMP^low^ and MMP^high^ cells. Size of the dots represent percent expression. (K) Representative UMAP plot shows gene expression-based cell fate bias of each type-II HSC cluster using Cell Radar analysis. Each cluster is color-coded with its respective blood cell type. (L) UMAP plot showing pseudotime trajectory analysis of pre-HSC type-I to type-II transition, with MMP subsets of the latter. Differentiation trajectory begins at pre-HSC type-I, with a red dot indicating the node.

To understand whether and how mitochondrial activity leads to transcriptional segregation of the definitive HSCs, UMAP-based clustering was performed on the pre-HSC type-II dataset, which revealed the presence of 8 transcriptionally distinct clusters C0-C7 (Figure 6E). Amongst these clusters, C2, C3, C4, C5, and C7 belonged to the MMP^low^ pool, and C0, C1, and C6 belonged to the MMP^high^ pool (Figure 6F). The expression levels of marker genes of each type-II HSC cluster showed striking differences between the MMP subsets, indicating the underlying transcriptional diversity (Figure 6G). As expected based on our functional assays, cell cycle analysis showed that the MMP^high^ pool was more quiescent, with a greater proportion of cells in the G1 phase and fewer in the G2/M phase, as compared to the MMP^low^ pool (Figure 6H, S6C). Further, the MMP^high^ pool showed increased expression of genes involved in oxidative phosphorylation and fatty acid metabolism (Figure 6I-J), with reduced expression of genes involved in mitochondrial quality control compared to MMP^low^ (Figure S6D). This is in agreement with the higher metabolic activity in MMP^high^ cells.

To uncover the identity of the cell clusters within the MMP subsets of type-II HSCs, fate mapping was performed to predict the blood cell bias. The marker gene set of each cluster (Table S4) was aligned with gene signatures of mature blood cell types using Cell Radar^13^ database to predict the cell fate bias. This showed that clusters belonging to the MMP^high^ subset are lymphoid biased, with C0, C1, and C6 exhibiting T cell, B cell, and lymphoid progenitor-like signatures, respectively. Clusters belonging to the MMP^low^ subset, however, showed myeloid fate bias with LT-HSC signature in C5, erythro-myeloid progenitor signatures in C2 and C4, and granulocyte and monocyte signatures in C3 and C7, respectively (Figure 6K; Figure S6E). Further, *asrij* was upregulated in the MMP^high^ subset (Figure S6F). This is in agreement with the *asrij* KO phenotype, where there is an expansion of the MMP^low^ subset in the AGM (Figure 5K) and a myeloproliferative disorder develops in the adult^18^, further strengthening our claim.

Pseudotime analysis of the embryonic HSC differentiation trajectory using Monocle 3 indicated that pre-HSC type-I transitions into the MMP^low^ pool, which in turn gives rise to the MMP^high^ pool (Figure 6L). Taken together, our results suggest that mitochondrial activity levels determine HSC fate during EHT, the earliest stage of definitive hematopoiesis. Further, MMP can potentially influence early HSC lineage segregation in the AGM itself.

### Mitochondrial activity modulates PI3K signaling to regulate embryonic HSC heterogeneity

To gain mechanistic insights into how mitochondrial activity regulates the differentiation of embryonic HSCs, the scRNA seq dataset of pre-HSC-type-II MMP subsets was used to perform a pseudo-bulk analysis. Raw read counts of each subset were acquired, and bulk-RNA transcriptome analysis pipeline was followed. PCA analysis revealed an 84% variance between the MMP subsets (Figure S7A), indicating a significant transcriptional separation, as inferred from our scRNA seq cluster analysis. DESeq2 analysis revealed 2036 DEGs (p-value < 0.05, log2FC >1 or <-1). MA plot (Figure S7B) and volcano plots (Figure 7A) represent the set of DEGs in the MMP^high^ subset compared to MMP^low^ (Table S5). A heatmap of all DEGs indicates enrichment of specific gene clusters across the two cell types (Figure 7B).

**Figure 7.**
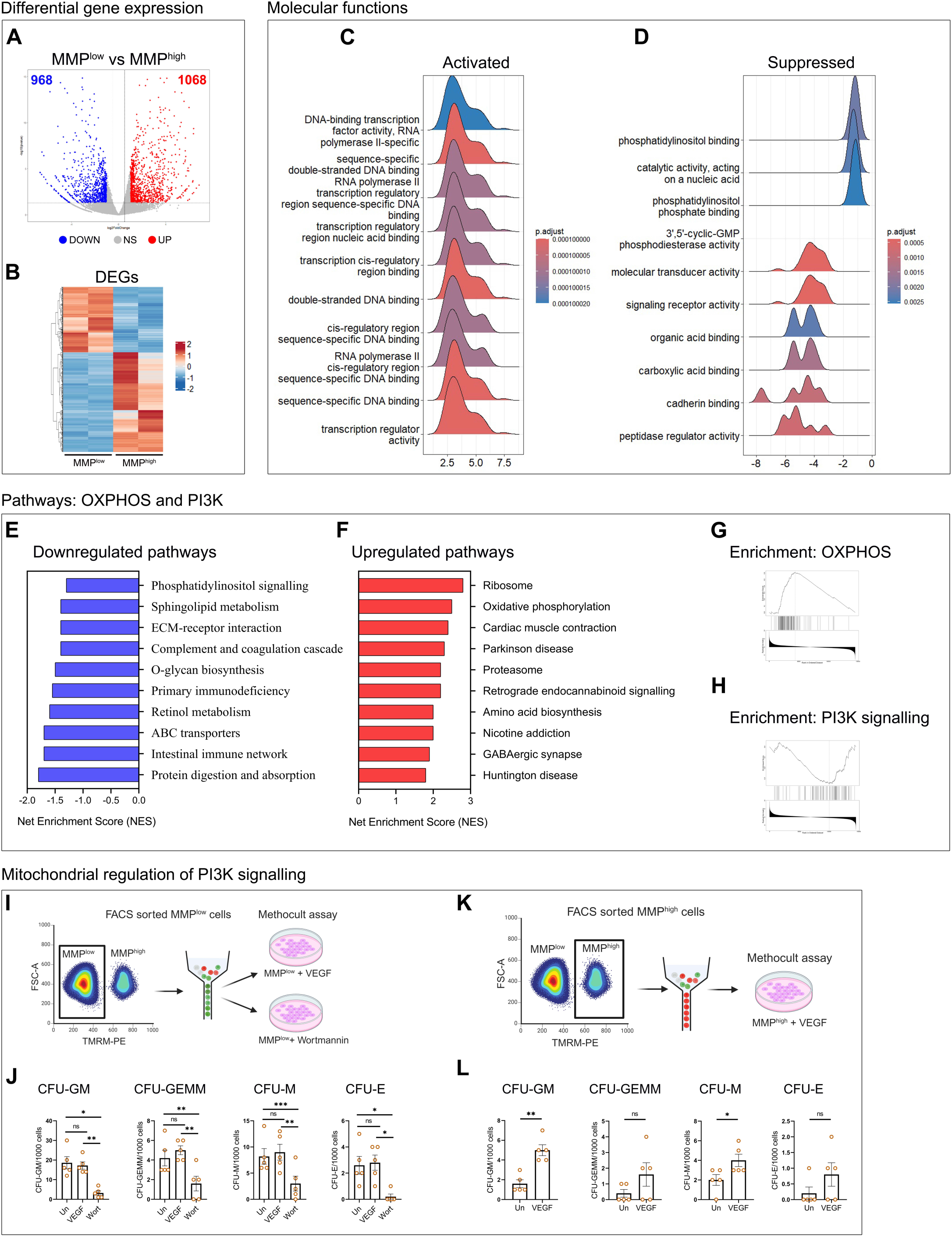
Mitochondrial activity modulates PI3K signaling to regulate embryonic HSC heterogeneity. (A) Volcano plot shows differentially expressed genes (DEGs) in WT E11.5 pre-HSC type-II MMP^high^ cells compared to MMP^low^ (red = upregulated, blue = downregulated, grey = non-significant; fold change > 1 or <-1, P value < 0.05) N = 2 independent experiments. See Table S5 for the gene list. (B) Heatmap representing all significant DEGs in WT E11.5 pre-HSC type-II MMP^high^ cells compared to MMP^low^. (C-D) Ridge plots represent molecular functions significantly activated (C) or repressed (D) MMP^high^ cells compared to MMP^low^. (E-F) Plots show net enrichment scores for signaling pathways upregulated (E) or downregulated (F) in MMP^high^ cells compared to MMP^low^. (G-H) GSEA enrichment plots for OXPHOS (G) and PI3K signaling (H) show enrichment scores or rank across the ordered dataset in MMP^high^ cells compared to MMP^low^. (I) Schematic representing FACS sorting of MMP^low^ subset followed by methocult assay in the presence of VEGF and Wortmannin. (J) The graph shows quantification of methylcellulose CFUs in untreated (Un), VEGF, or Wortmannin (Wort) treated MMP^low^ subset of WT E11.5 pre-HSC type-II. N = 5, n = 1000 HSCs from ≥ 7 pooled AGM, data represent mean ± SEM. (K) Schematic representing FACS sorting of MMP^high^ subset followed by methocult assay in the presence of VEGF. (L) Graph shows quantification of methylcellulose CFUs in untreated (Un) and VEGF-treated MMP^high^ subset of WT E11.5 pre-HSC type-II. N = 5, n = 1000 HSCs from ≥ 7 pooled AGM, data represent mean ± SEM.

At the molecular level, transcription factor binding and RNA polymerase-II activity was upregulated in the MMP^high^ subset, while molecular functions like phosphatidyl inositol binding, receptor binding and catalysis was downregulated in MMP^high^ cells (Figure 7C-D). Further, gene set enrichment analysis (GSEA) was done to identify the cellular pathways and processes involved. PI3K signaling, sphingolipid metabolism, and extracellular matrix (ECM) interactions were downregulated, with an upregulation of oxidative phosphorylation (OXPHOS), ribosome biogenesis, and amino acid biosynthesis in the MMP^high^ subset (Figure 7E-F). Enrichment plots for oxidative phosphorylation and PI3K signaling also reflect significant upregulation and downregulation, respectively, across the MMP^high^ gene sets (Figure 7G-H). Along with this, biological processes like blood vessel morphogenesis, angiogenesis, and vascular development were significantly downregulated in the MMP^high^ subset (Figure S7C), processes that are known to be regulated by Vascular endothelial growth factor (VEGF)-mediated PI3K signaling^43^.

Several PI3K signaling target genes were downregulated in the MMP^high^ subset (Figure S7D; Table S6). To further investigate the regulatory axis governing the differentiation potential of MMP subsets, PI3K signaling was modulated in the MMP^low^ and MMP^high^ type-II HSC pool, followed by assaying for HSC differentiation potential. First, the MMP^low^ pool was FACS-sorted and cultured in methylcellulose medium in the presence of VEGF, an activator of PI3K signaling^44^, or with Wortmannin, a well-established inhibitor of PI3K signaling^45^ (Figure 7I). The differentiation potential of MMP^low^ HSCs under PI3K modulation was compared to the vehicle treated control. There was no change in the number of CFUs in VEGF treated cells compared to control, possibly owing to already high PI3K signaling in MMP^low^ cells. However, a lower number of CFUs were observed in Wortmannin treated MMP^low^ cells, resembling the differentiation phenotype of the MMP^high^ subset (Figure 7J). Conversely, MMP^high^ cells were cultured in the presence of VEGF to upregulate the PI3K signaling in this subset (Figure 7K). Again, we observed a reversal of the differentiation phenotype with a greater number of CFU-G and CFU-M in VEGF treated samples compared to controls (Figure 7l). Thus, our results indicate that reduced mitochondrial activity results in upregulation of the PI3K signaling cascade, thereby increasing differentiation in the mature embryonic HSCs.

## Discussion

Single-cell transcriptomic studies have highlighted the inherent heterogeneity of gene expression in HSC populations, and elegant *in-vitro* and *in-vivo* experiments have correlated this with their potency^24,25,31,38,46–48^. However, the isolation of sufficient numbers of desired populations of repopulating HSCs based on gene expression is confounding and challenging^49,50^. Further, efforts to generate definitive HSCs by *in-vitro* differentiation have had limited success^51–53^. Recent studies showed that bone marrow HSC potency is governed by mitochondrial status^13,14,41^. Further, metabolic regulation can help reduce or reverse HSC aging, giving rise to youthful HSCs with improved repopulation capacity^13^. However, there is a major gap in the understanding of how and when metabolic heterogeneity in HSC arises and whether it is innate or an outcome of the myriad extrinsic influences on the adult HSC.

In this study, we trace the earliest origins of HSC heterogeneity to the emerging HSCs in the embryonic AGM *in-vivo*. We show that mitochondrial activity is a key determinant of HSC potency and lineage choice. We identify that lympho-myeloid fate choice is first made in the mature AGM HSCs, and depends on mitochondrial membrane potential. Further, by genetic and pharmacological means, we demonstrate that modulating mitochondrial activity or associated signaling pathways during the endothelial to hematopoietic transition can change HSC emergence and phenotype in a predictable manner. Therefore, fine-tuning mitochondrial activity during specific developmental windows could enable robust production of definitive HSCs.

An increase in mitochondrial respiration in human pluripotent stem cells (hPSCs) was shown to induce definitive hematopoiesis *in-vitro*^33^. Further, *in-vitro* human EHT was associated with a gradual increase in mitochondrial activity^33^. In mouse embryonic development, the first events of definitive hematopoiesis require the hemogenic endothelial cells to undergo a shape change and delaminate as spherical HSC (type I) that mature and gain repopulation potential (type II)^54^. It is reasonable to assume that this energy-intensive process would require a significant increase in mitochondrial mass, which we observed in the emerging AGM HSCs. Drp1-mediated fission was increased during HSC emergence, giving rise to short, seemingly immature mitochondria, generally associated with quiescent cells. However, the increase in mitochondrial activity, ROS, and mitophagy are indicative of metabolic stress in emerging HSCs, which was unexpected for cells that should become quiescent^14,42^. The transition from emerging to mature HSCs, however, was accompanied by a decrease in mitochondrial content, ROS, and calcium in type-II HSCs. This suggests that the energy burst seen in emerging HSCs is transient and must be reset to baseline mitochondrial activity, possibly to reduce the cellular damage in the long-term HSCs that will populate the bone marrow. Indeed, the majority of type II HSCs showed low MMP. However, there was a gradient of MMP, and a subset of the type II HSC pool had several folds higher MMP, skewing the total MMP of the population to a higher level. Thus, newly emerged definitive HSCs in the AGM are a heterogeneous population that can be segregated based on their mitochondrial membrane potential. Our study provides the first evidence of mitochondrial heterogeneity in embryonic definitive HSCs

Previous studies show that an increase in mitochondrial respiration results in the production of fewer intra-aortic hematopoietic clusters (IAHCs). However, this does not help distinguish between the impact on HSC emergence and maturation. Our findings that mitochondrial parameters differ between type-I and type-II HSCs raise the possibility that mitochondria may affect each stage of EHT differently. Indeed, while lowering mitochondrial activity in the AGM did not affect HE and type-I HSC frequency, there was a significant expansion of type-II HSCs. This was observed with pharmacological inhibition of mitochondrial activity with CCCP *ex-vivo* as well as in the *asrij* KO model *in-vivo*. Increased HSC numbers could be an outcome of enhanced type-I to type-II HSC differentiation, or increased proliferation of type-II HSCs. Our bulk transcriptome analysis indicated that cell proliferation signatures were unchanged. Interestingly, Wnt signaling, which is known to promote definitive hematopoiesis in the embryo^55–58^ via cell fate transition, was upregulated. Thus, it is likely that reducing mitochondrial activity in the transiting HSC pool accelerates HSC maturation by upregulation of Wnt signaling, thereby increasing type-II HSC frequency. We, therefore, establish a cause-and-effect relationship between mitochondrial activity levels and the development of definitive embryonic HSCs during EHT. Hence, our study opens up new opportunities for understanding the earliest stages of hematopoiesis *in-vivo*.

Definitive HSCs of different potencies are experimentally isolated by the use of surface marker combinations or reporter gene expression. However, this approach is limited by the availability of functional correlates of gene expression. Transcriptional profiling allows in-depth analysis of gene expression but does not retain cell integrity. In contrast, HSCs can be sorted based on their MMP status using dyes/probes that are agnostic to gene expression status and do not compromise cell viability and potential. Further, the functional relevance of mitochondrial heterogeneity is highlighted by studies demonstrating the repopulating ability of HSCs with varying MMP and reversibility by metabolic manipulation^13^. Taking advantage of these attributes, we isolated pre-HSC type-II based on their MMP level and showed that they differ in their potential both *in-vitro* and *in-vivo*. While low mitochondrial activity promotes enhanced differentiation, high mitochondrial activity induces HSC quiescence during embryonic stages. In addition, this is corroborated by the *asrij* KO AGM, where an expansion of the MMP^low^ pool leads to enhanced differentiation, which is sustained throughout adult hematopoiesis.

AGM HSCs undergo the next stage of development in the fetal liver, where they expand to give rise to hematopoietic stem and progenitor cells (HSPCs)^30,59–61^. Fetal hematopoiesis shares many similarities with the hematopoietic hierarchy in the BM, with LT-HSCs giving rise to the downstream multipotent progenitors. While BM LT-HSCs show an equal distribution of MMP^low^ and MMP^high^ subsets, the profile for FL LT-HSCs has not been reported. Our analysis indicates that though AGM type-II HSCs are skewed towards the MMP^low^ pool, the distribution is balanced by the time they populate the fetal liver. However, whether this matches the potency of the bone marrow MMP subsets remains to be tested.

Apart from fate determination, mitochondrial activity levels also determine HSC lineage bias. Levels of mitochondrial tricarboxylic acid cycle (TCA) metabolites like glutamine and pyruvate in human HSCs generated through *in-vitro* EHT can determine HSPC lineage bias ^32,33^. In mouse BM LT-HSCs, mitochondrial activity levels affect lineage choice. While myeloid or platelet-biased genes were upregulated in MMP^low^ HSCs, genes involved in lymphoid and erythroid priming were upregulated in MMP^high^ HSCs^13^. Our single-cell analysis of E11.5 HSCs corroborates the impact of MMP levels on lineage choice. While MMP^low^ AGM HSCs show transcriptional signatures of myeloid cells, MMP^high^ HSCs are lymphoid-biased. Expansion of the MMP^low^ subset in *asrij* KO AGM correlates with a myeloproliferative disorder in the adult BM^18^. Thus, metabolically segregated HSCs show that lineage bias arises in the AGM itself. However, when the lineages segregate is not known. Our trajectory analysis suggests that the more abundant myeloid-biased HSCs give rise to the lymphoid-biased HSCs.

Interestingly, a recent study reported two separate populations of CD45^+^ HSPCs at E10.5, reflecting an initial wave of lymphomyeloid-biased progenitors, followed by HSC precursors^62^. Therefore, subsets of the CD45^+^ cells reported previously could be driven by mitochondrial activity, where the lymphoid-biased definitive HSCs at E11.5 could be derived from the HSC precursors found at E10.5. Further, our scRNA seq analysis reveals a distinct set of genes that mark the MMP^low^ and MMP^high^ subsets. Genetic barcoding or fluorescent tagging of a few candidate genes, followed by *in-vivo* tracking during hematopoietic development, could help establish novel signatures of clonal lineage-biased HSC subsets.

Mechanistically, we found that lower mitochondrial activity in HSCs upregulates the PI3K signaling pathway to promote differentiation. While PI3K signaling is known to modulate HSPC stemness in zebrafish^63^, its regulation in mammalian embryonic hematopoiesis is not clear. We show that modulation of PI3K led to a reversal of the differentiation potential in the MMP^low^ or MMP^high^ subsets. The presence of elevated levels of PI3K signalling in the *asrij* KO BM^18^ support this result. Thus, metabolic segregation of the HSC pool translates to differential PI3K signaling for HSC fate determination and lineage choice.

In summary, our study provides a detailed analysis of how mitochondrial metabolic tuning in the developing HSCs orchestrates the cellular transcriptome and signaling pathways to regulate HSC potential during embryogenesis.

## Limitations of the study and future directions

Our study explores how mitochondrial activity modulates cellular signaling pathways to regulate the potency of early embryonic HSCs. However, we have used MMP as a holistic readout of mitochondrial activity. Since there are multiple factors that govern MMP itself, this study opens up the possibility of examining the contribution of each of these factors. For instance, it would be interesting to understand the impact of the electron transport chain complexes using specific inhibitors against each complex and assaying for HSC frequency and function. Further, the *asrij* KO model used in our study is a global KO. Using a conditional KO to deplete *asrij* in subpopulations of the AGM during EHT will help identify the cell-autonomous effect of mitochondrial modulation. Also, additional genetic models such as *Drp1* and *Mfn2* conditional KO mice can be used to understand how fission-fusion dynamics affect EHT, while *Tfam* conditional KO can be used to study the impact of mitochondrial biogenesis. Overall, for most biochemical and metabolic assays, the extremely rare and transient nature of these embryonic HSCs, and the inability to grow or expand them *in-vitro* under niche-free conditions, precludes some of these analyses.

## Methods

### Mouse models

C57BL/6J (#:000664) mice were purchased from the Jackson laboratory (Bar Harbor, Maine, USA). The generation of and genotyping strategy for the R26-Mito-EGFP (#CDB0251K) strain has been described previously^64^ and was obtained from Riken. The R26R-mito-EGFP mice were crossed with PGKCre (JAX #020811) to generate the R26-Mito-EGFP, Tg-PGKCre. *asrij* floxed and KO mice generation and genotyping has been described previously^18^. All mice strains were maintained and bred in the JNCASR animal facility in individual cages under a 12-hour light-dark cycle, with a chow diet and water ad libitum. All mice experiments were performed in accordance with the Institutional Animal Ethics Committee (IAEC) of JNCASR. See Table S1 for genotyping primer details.

### Mouse embryo dissection and tissue isolation

Timed mating was performed using young mice (8-16 weeks) of the desired strain to obtain mid-gestation embryos. Observation of the vaginal plug was considered as E0.5 of embryonic development. At the desired time point, pregnant dams were euthanized, and the embryos were isolated from the uterine horn. Precise staging of the embryo was done on the basis of vaginal plug date and embryo somite numbers. AGM region was dissected from E9.5 to E11.5 embryos. The fetal liver was isolated from E15.5 embryos. For BM isolation, 2-4-month-old mice were euthanized, followed by dissection of the femur and tibia from both hind limbs. Bone marrow cells were isolated by flushing the bones in 1X phosphate buffer saline (PBS) with 0.05% sodium azide. The cells were pelleted at 3000 rcf for 3 minutes, followed by resuspension in 1X PBS.

### Embryo cryosectioning

Embryos of the desired stage were dissected and fixed in 4% paraformaldehyde (PFA) (#GRM3660, HiMedia) for 1 hour at room temperature (RT), followed by washing with 1X PBS. Successive rounds of tissue equilibration were done overnight with 15% and 30% sucrose (#28105, Thermo Fisher Scientific) at 4°C. Equilibrated embryos were embedded and frozen in PolyFreeze Tissue Freezing media (#SHH0026, Sigma-Aldrich) and stored at -80°C until used for sectioning. 20 µm sagittal sections were taken on a cryostat (#CM3050S, Leica) across the dorso-ventral axis of the embryo, and collected on 0.3% gelatin (#D12054, Invitrogen) coated slides.

### Immunofluorescence staining

Sections were washed with 1X PBS, followed by permeabilization with 0.3% Triton X-100 (#T8787, Sigma-Aldrich) in 1X PBS for 30 minutes at RT. The sections were washed twice with blocking solution (4% fetal bovine serum (FBS) in 1X PBS) for 15 minutes each, then incubated in blocking solution for 1 hour at RT, followed by incubation with the desired primary antibodies overnight at 4°C. Primary antibodies were used in the following dilutions: c-kit (1:100; # AF1356, R&D Biosystems), CD31 (1:50; #553370, BD Biosciences), Tom20 (1:100; #11802-1-AP, Proteintech), Drp1 (1:50; #ab184247, Abcam), pDrp1^S616^ (1:50; #3455, Cell Signalling Technology), LAMP1 (1:100; #ab24170, Abcam). All secondary antibody incubations were performed for 1 hour at RT, at 1:400 dilution: Alexa fluor (AF) 488, 568, or 633 of anti-goat, anti-rat, and anti-rabbit antibodies (Invitrogen). Post secondary antibody staining, sections were washed twice with 1X PBS and mounted in 4′,6-diamidino-2-phenylindole (DAPI) (1:400) to stain the nucleus, along with ProLong Gold Antifade Mountant (#P36930, Invitrogen) in 1:1 ratio.

### Confocal imaging and analysis

Images were acquired on a Zeiss LSM 880 confocal microscope at 20X, 40X, and 63X magnification with 0.5 μm *z*-interval. Airy scan mode was used at 63X zoom 3, at 0.3 μm *z*-interval for high-resolution imaging. Image analysis and processing was done using ImageJ software. For analysis of median fluorescence intensity (MFI), the region of interest (ROI) was marked using the freehand selection tool of ImageJ, and fluorescence intensity normalized to the area was measured. Specifically, for analysis of mitochondrial mass across HE and IAHCs, each c-kit^+^ cell of the IAHC was selected for HSC analysis, along with the selection of the underlying endothelial cell for comparison. Analysis of the mitochondrial network was done using the Mitochondrial Network Analysis (MiNA) plugin of the Fiji software according to a previously published protocol^65^. 3D reconstruction of the mitochondrial surface was made using the surface tool of Imaris software. Mito-Drp1 and Mito-LAMP1 analysis was done using the spot-surface interaction algorithm of the Imaris software. Total spots, spots in contact with the surface, and spots per unit surface area were calculated using 0.3 μm as the distance threshold across both HE and HSCs. The surface volume function was used to calculate the volume of mitochondrial surface across HE and HSC. Each embryo was considered as a biological replicate and 25-30 cells of each HE and HSC were analysed per replicate.

### Fluorescence-activated cell sorting (FACS) of embryonic, fetal, and BM HSCs

Post dissection, AGM tissue dissociation was done by incubating with 0.2% collagenase (#17104-019, Gibco) at 37°C for 15 minutes, followed by gentle pipetting and then passed through a 70 µm cell strainer (#352350, Falcon) to obtain a single cell suspension in 1X PBS and 0.05% sodium azide. For analysis of the embryonic HE and HSC pool, resuspended AGM cells were incubated with surface markers c-kit-APC (#105812, BioLegend) or c-kit-PE (#12-1171-82, eBiosciences), CD31-PECy7 (#102524, BioLegend) and CD45-FITC (#103108, BioLegend) or CD45-APC-Cy7 (#103116, BioLegend) for 1 hour at 4°C. All antibodies were used at 1:100 dilution. Cells were washed and resuspended in 1X PBS for analysis/sorting. Fetal liver cells were dissociated using gentle pipetting, and clumps were removed until a single-cell suspension was obtained. Cells were resuspended in 1X PBS with 0.05% sodium azide. For fetal liver or BM HSPC analysis, cells were depleted for lineage-committed progenitors either using the Lin cocktail-APC (#560492, BD Biosciences) or using the Lin cocktail beads (#130-110-470, Miltenyi Biotech) and magnetic separation (MACS) according to manufacturer’s instructions. Post depletion of lineage committed progenitors, cells were stained with surface markers-Sca1-PECy7 (#561021, BD Biosciences), c-kit-APC-Cy7, CD150-FITC (#11-1502-82, eBiosciences), and CD48-BV421 (#103428, BioLegend) or CD48-APC (#17-0481-82, eBiosciences) for 1 hour at 4°C. Cells were washed with 1X PBS post staining and prior to FACS sorting or analysis, 7-Aminoactinomycin D (7AAD) (#559925, BD Biosciences) was added to allow dead cell exclusion. For intracellular staining, cells were fixed with 4% PFA for 10 min at 4°C post surface staining. This was followed by 1X PBS wash and permeabilization using 0.1% triton-X for 10 minutes at 4°C. Cells were then incubated overnight with the following primary antibodies-Tom20 (1:100; #11802-1-AP, Proteintech), Drp1 (1:100; #ab184247, Abcam), p-Drp1^S616^ (1:100; #3455, Cell Signalling Technology). Secondary antibody incubation was then done for 1 hour at 4°C with Alexa fluor 488/568/633 (Invitrogen) at 1:400 dilution of the desired host. Unstained cells, isotype control, and fluorescence minus one (FMO) control were used in all experiments. All samples were analysed on BD FACS Aria-III (BD Biosciences) or BD FACS Fusion (BD Biosciences), with the BD FACS Diva software. Analysis of the data was performed using FlowJo software v10.8.1 (BD Biosciences).

### Flow cytometry analysis of mitochondrial activity, lysosomal content, and glucose uptake

After staining for surface markers of the desired cell type, the cell suspension was washed, and resuspended in 1X PBS with either of the following mitochondrial probes for 15 minutes at 37°C: MitoTracker Deep Red (25nM; #M22426, Invitrogen), Tetramethylrhodamine, Methyl Ester, Perchlorate (TMRM) (100nM; #T668, Invitrogen), MitoSOX Red (5μM; #M36007, Invitrogen). For analysis of lysosomal content, cells were incubated with LysoTracker Deep Red (25nM, #L12492, Invitrogen). For glucose uptake analysis, cells were incubated with 2-NBDG (200μM; N13195, Invitrogen) for 15 minutes at 37°C. Post staining cells were washed with 1X PBS and used for flow cytometry analysis. Unstained cells, isotype control, and FMO controls were used in all experiments.

### Reverse transcription-quantitative PCR (RT-qPCR)

HE or HSCs were FACS-sorted in 200μL TRIzol reagent (# 15596026, Invitrogen) and lysed by vortexing. RNA isolation was done from the lysate as per the manufacturer’s instructions. cDNA synthesis was done through reverse transcription using the SuperScript III first-strand synthesis kit (#18080051, Invitrogen) following the manufacturer’s instructions. RT-qPCR was set using SensiFAST SYBR No-ROX (#BIO-98020, Bioline) in CFX384 real-time PCR system (Bio-Rad) as per the manufacturer’s protocol. Primer sequences used are mentioned in Table S1. Gene expression was normalized to *Beta-2 microglobulin* (*B2m)* expression.

### ATP assay

Intracellular ATP was measured using the luminescent ATP detection assay kit (#ab113849, Abcam) according to the manufacturer’s instructions. Briefly, FACS-sorted cells were lysed by adding 50 μL detergent, followed by incubation in an orbital shaker at 600 rpm for 5 minutes. 50 μL substrate was then added and incubated in the shaker at 600 rpm for 5 minutes. The samples were then incubated at RT for 10 minutes, and luminescence was recorded using the Varioskan™ LUX multimode microplate reader (#VL0000D0, Thermo Scientific).

### AGM explant culture assay

AGM tissue from E10.5 and E11.5 embryos was dissected and cultured on a 0.65 μm membrane filter (#DAWP04700, Sigma) at the gas-liquid interface for 48 hours in center well organ culture dishes (#353037, BD Falcon), with 1.5 mL MyeloCult media (#H5100, Stem Cell Technologies) supplemented with 10^-6^M hydrocortisone (#H2270, Sigma) and 1X antibiotic-antimycotic (#15240-062, Invitrogen). The media was supplemented with either CCCP (5 μM, #C2759, Sigma) or Mitoquinol (7 μM; #89950, Cayman chemical company) or IWP4 (5 μM, #SML1114, Sigma). For lineage marker analysis, tissues were kept under culture for 72 hours. Post treatment, AGM dissociation was done using 0.2% collagenase for 15 min at 37°C and passed through a 70 μm cell strainer to obtain a single cell suspension. The cell suspension was stained with the desired cell surface markers and analyzed through flow cytometry.

### *In-vivo* CCCP treatment

Pregnant dams were injected with CCCP 24 hours prior to analysis. CCCP was administered through intraperitoneal injection at 0.5 mg/kg body weight, and dilutions were made in 100 μL 1X PBS. Control mice were injected with 100 μL PBS.

### Methylcellulose culture assay

MMP subsets of pre-HSC type-II were sorted in equal numbers in 100 µl Iscove’s modified Dulbecco’s medium (IMDM) media (#12440053, Gibco) supplemented with 2% FBS. Cell suspension was added to 1.5 mL methylcellulose media (MethoCult GF #M3434, Stem Cell Technologies) supplemented with 1X antibiotic-antimycotic (#15240-062, Invitrogen) and plated on 35 mm petri dishes (#351008, BD Falcon). In the case of treatments, media was supplemented with modulators of PI3K signaling-VEGF (50 ng/mL; #V7259, Sigma-Aldrich) or Wortmannin (5 µM; #95455, Sigma-Aldrich). The cells were cultured at 37°C, and colonies were scored after two weeks.

### Hematopoietic and endothelial dual potential assay using OP9 co-culture

OP9 cells were acquired from American Type Culture Collection (ATCC) (#CRL-2749) and cultured in Alpha Minimum Essential Medium (α-MEM #12561056, Gibco) supplemented with 20% FBS, 2.2 g/L sodium bicarbonate (#S5761, Sigma) 1% GlutaMAX (#35050061, Invitrogen) and 0.05% β-mercaptoethanol (#M7522, Sigma) on 0.1% gelatin-coated wells. Cells were passaged using 0.25% (w/v) Trypsin-0.53 mM EDTA solution (#15400-054, Invitrogen). FACS-sorted AGM HSCs were seeded on a 40-50% confluent layer of OP9 cells in an OP9 co-culture medium composed of IMDM, 2% FBS, 50 ng/mL VEGF, 50 ng/mL SCF (#250-03, Peprotech), 100 ng/mL IL-3 (#I1646, Sigma) and 100 ng/mL Flt3 ligand (#F3422, Sigma). After 14 days of co-culture, cells were harvested through vigorous pipetting and stained for surface markers of the desired cell lineages: endothelial (CD31-PECy7), pan-hematopoietic (CD45-FITC), myeloid (CD11b-FITC), B cells (CD19-FITC, B220-APC), T cells (CD3-FITC), and HSPCs (Lin-APC, c-kit-PE, Sca1-PECy7). All antibodies were used at 1:100 dilution and cells were analyzed through flow cytometry.

### Hematopoietic stem cell (HSC) transplantation assay

The recipient mice (C57BL/6J) were fed acidified water (pH 1.8-2.6) with Enrofloxacin (0.16 mg/mL) for one week before irradiation, and lethally irradiated at 9 Gy (1.3 Gy/min). 5 X 10^5^ unfractionated BM cells from C57BL/6J mice were mixed with FACS-sorted MMP^low^ or MMP^high^pre-HSC type-II cells from R26-mito-EGFP mice in 100 µL IMDM with 2% FBS. Cell suspension was injected into the lethally irradiated recipients via retro-orbital injection. Recipient mice were fed acidified water for up to 10 days after irradiation. Peripheral blood was drawn via retro-orbital bleeds for flow cytometry analysis performed at 1-month intervals for 4 months post-transplantation. At the end of 4 months, BM was harvested and stained for HSPCs (Lin-APC, c-kit-PE, Sca1-PECy7), B cells (CD19-PECy7), T cells (CD4-APC, CD8-PE), myeloid cells (CD11b-APC) and pan hematopoietic cells (CD45-APC-Cy7). The percentage of GFP-positive cells was analyzed to calculate the degree of chimerism in both peripheral blood and bone marrow.

### Bulk RNA transcriptome analysis

E10.5 control and CCCP-treated HSCs were MACS-sorted from AGM explants using c-kit microbeads (#130-091-224, Miltenyi Biotech). RNA was isolated through the TRIzol method and quantified using a Qubit high-sensitivity assay kit (#Q32852, Invitrogen) on a Qubit 4 fluorimeter (Invitrogen). Samples were run on the TapeStation system (Agilent) with high-sensitivity RNA Screen Tape analysis (Agilent, #5067-5579) for RNA quality control (QC). cDNA preparation, library construction, and RNA sequencing were performed by the Next Generation Sequencing (NGS) facility at the Bangalore Life Science Cluster (BLiSC), Bengaluru, India. Briefly, mRNA isolation was done using the NEBNext Poly(A) mRNA Magnetic Isolation Module (#E7490L). cDNA stranded library construction was performed using NEBNext® Ultra II Directional RNA Library Prep with Sample Purification Beads (#E7765L) according to the manufacturer’s instructions. 30 million reads were sequenced on the Novaseq6000 platform using 2×100bp sequencing read length.

Quality check of the sequenced FASTQ files was done using the FastQC tool, followed by trimming, which involves the removal of adaptor sequences and low-quality bases (Q < 20) from the ends of the reads using Fastp. UCSC mm10 was used as the reference genome and indexed using Hisat2. Trimmed reads were aligned to the indexed genome using Hisat2. The resulting Sequence Alignment/Map (SAM) files were sorted by coordinates and converted to Binary Alignment/Map (BAM) files using Samtools. Gene expression values were calculated using the Feature counts package of the Subread R package, and summarised counts for each gene were obtained. This file was further used for differential gene expression analysis using the DESeq2 package in R. Differentially expressed genes (DEGs) with log2FC > 0.5 and p value < 0.05 were considered upregulated and log2FC < 0.5, p value < 0.05 were considered downregulated. All plots were generated using the ggplot package. Heatmaps were generated using pheatmap or complex heatmap packages. Cluster Profiler package was used for gene set enrichment analysis and gene ontology analysis. Pathway enrichment analysis was done using the WEB-based Gene Set Analysis Toolkit, which has integrated datasets from KEGG, Panther, Reactome, and Wiki pathways.

### Single-cell RNA sequencing and analysis

Freshly sorted CD31^+^ AGM cells based on TMRM intensity-MMP^low^ and MMP^high^ were resuspended in 1X PBS with 2% FBS. Cell suspension was loaded into the Single Cell B Chip (10X Genomics, USA) and processed in the Chromium single cell controller (10X Genomics, USA), targeting a maximum of 10,000 cells per lane. The single-cell mRNA library was prepared using the Chromium Single Cell 3’ Reagent kit v3 (10X Genomics, USA) according to the manufacturer’s instructions. Sequencing was performed on the Novaseq6000 platform using an SP-V1.5-100 cycle kit with a read depth of 300-400 million reads. Two independent biological replicates were performed and an entire litter of 7-9 embryos was used per experiment for FACS. The cell ranger 2.1.1 pipeline (10X Genomics) was used to align reads to the mm10 reference genome, and barcode matrices were generated using the default parameters.

All downstream processing and data analysis was done using the Seurat v4.0.6 pipeline. Pre-processing and quality control was done for the removal of cells with read counts greater than 2,500 or less than 200 to remove doublets or dead cells, respectively. Additionally, cells with > 20% genes mapping to the mitochondrial genome were also excluded from the analysis. This was followed by log normalization using the NormaliseData function, identification of highly variable features using the Vst function, and data scaling using the ScaleData function. The scaled data was further used for linear dimensionality reduction and PCA analysis using the RunPCA function. Cell clustering was performed based on the nearest neighbor algorithm using the FindCluster function, using 0.5 resolution and the first 10 dimensions. Non-linear dimensional reduction was performed using the runUMAP function. Differentially expressed features or cluster biomarkers were determined using the FindAllMarkers function, followed by assigning cell type identity to the clusters. FeaturePlot, DoHeatMap, and Dotplot functions were used respectively to plot feature plots, heatmaps, and dotplots.

### Cell cycle, pseudotime trajectory analysis, and pseudobulk analysis

Cell cycle analysis was performed using Seurat’s CellCycleScoring function based on the expression of canonical markers. The function identifies cells based on their levels of G2/M and S phase markers.

Pseudotime analysis was performed using the Monocle 3 package. The expression matrix of the Cell Ranger output file was stored in a cell data set (cds) object, which was further used for batch correction and clustering using the align_cds and cluster_cell functions, respectively. Cells were ordered and plotted in a pseudotime trajectory using the learn_graph and order_cells function.

For pseudobulk analysis, raw UMI counts of the desired cell type were pooled using the read_counts function and passed through the bulk-RNA seq analysis pipeline as mentioned above.

### Statistical analysis

All data are represented as mean ± Standard error of mean (SEM). Statistical significance was calculated using one of the following tests based on the experiment: one sample t-test (for experiments involving fold change analysis), unpaired Student’s t-test (for comparison between two groups), and one-way analysis of variance (ANOVA) with Tukey’s multiple comparisons post-hoc test (for comparison between more than two groups), and paired Student’s t-test (for HSPC bone marrow chimerism analysis). A P-value less than 0.05 was considered significant, ns - not significant, *P<0.05, **P<0.01, ***P<0.001. Graphs were plotted using Prism 10 software (Graphpad).

### Data and code availability

Our study does not report any original codes. Bulk and scRNA seq datasets are accessible through GEO accession numbers GSE274537 (bulk RNA sequencing) and GSE274538 (scRNA sequencing). Any additional data required will be made available upon request.

## Acknowledgments

We thank members of the Inamdar laboratory and JNCASR community for their fruitful suggestions and valuable inputs; Prerana M for help with single-cell cDNA library preparation; Prathamesh Dongre for help with mice intraperitoneal injections; JNCASR bioimaging, flow cytometry, and animal facility; Department of Biotechnology, Government of India for funding. BioRender was used for schematics and graphical abstract.

## Authors’ contributions

Conceptualization: A.P and M.S.I; Methodology, experiments, and data analysis: A.P; Manuscript writing, review, and editing: A.P and M.S.I; Funding acquisition and resources: M.S.I; Supervision: M.S.I.

