## Supplementary text and figures for "Mitochondrial activity-driven hematopoietic stem cell fate and lympho-myeloid lineage choice is first established in the aorta-gonad-mesonephros"

**Supplementary information:**

Supplementary text and figures: Supplementary figures S1-S7, Table S1 and supplementary figure legends

Table S2-S6: Excel files containing bulk and scRNA-seq gene expression values

**Figure S1. HE to pre-HSC type-I transition is marked by an increase in mitochondrial activity and content.**

(A) Representative flow cytometry scatter plots show the gating strategy for WT E11.5 AGM hemogenic endothelial (HE) cells, pre-HSC type-I, and pre-HSC type-II.

(B-C) Flow cytometry histograms show EGFP intensity in HE (black) and HSC-I (blue) of E9.5 (B) and E10.5 (C) R26-mito-EGFP embryos. Graph shows fold change in EGFP MFI. N = 3, n ≥ 7 embryos, and data represent mean ± SEM.

(D-E) Flow cytometry histogram shows MitoTracker-Deep Red (D) and Tom20 (E) intensity in HE (black) and HSC-I (blue) of WT E11.5 embryos. Graph shows fold change in MFI. N = 4, n ≥ 7 embryos), and data represent mean ± SEM.

(F-G) Flow cytometry histogram shows TMRM intensity in HE (black) and HSC-I (blue) of WT E9.5 (F) and E10.5 (G) embryos. Graph shows fold change in TMRM MFI. N = 3, n ≥ 7 embryos, and data represent mean ± SEM.

**Figure S2. Heterogeneity in the expression of mitochondrial genes within definitive HSCs.**

(A) UMAP plots show expression of mitochondrial genes- *Opa1, Tomm20, Tfam, Cox4, Mfn1* and *Mfn2* in the E11.5 c-kit$^{+}$ Gata2$^{low}$ HSC pool, adapted from Vink et al., 2020[1].

(B-C) Representative flow cytometry histograms show TMRM intensity profile in HE, HSC-I and HSC-II in WT E10.5 (B) and E11.5 (C).

**Figure S3. Lower mitochondrial activity in the AGM leads to expansion of the pre-HSC type-II pool.**

(A-B) Representative flow cytometry scatter plots show frequency of pre-HSC type-II in untreated and CCCP treated WT AGM explants at E10.5 (A) and E11.5 (B).

(C-D) Representative flow cytometry scatter plots show frequency of pre-HSC type-II in untreated and CCCP treated WT pregnant dams at E10.5 (C) and E11.5 (D).

(E) Graph shows fold change in TMRM MFI in *asrij* KO compared to floxed in pre-HSC type-II. N = 6, n ≥ 7 embryos, and data represent mean ± SEM.

(F) Representative flow cytometry scatter plots show frequency of pre-HSC type-II in *asrij* floxed and KO AGM at E11.5.

**Figure S4. Lower mitochondrial activity upregulates Wnt signaling to promote pre-HSC type-II expansion.**

(A) Principal component analysis (PCA) of untreated and CCCP treated WT E10.5 AGM c-kit$^+$ HSC transcriptome. N=2 independent experiments.

(B) MA plot of CCCP vs untreated HSCs.

(C) Bar plot shows downregulated pathways upon CCCP treatment in WT E10.5 HSCs, highlighting the TGFβ signaling pathway.

(D) Heatmap shows expression of genes involved in the TGFβ signalling pathway in untreated and CCCP treated HSCs.

(E-F) Graphs show HE and HSC-I frequency in WT AGM explants upon CCCP or CCCP+IWP4 treatment at E10.5 (E) and E11.5 (F).

**Figure S5. Mitochondrial membrane potential based heterogeneity in AGM and fetal liver HSCs.**

(A) Representative flow cytometry scatter plots show TMRM intensity-based distribution of HE, pre-HSC type-I and pre-HSC type-II in WT E10.5 AGM. Graph shows frequency of $MMP^{low}$ and $MMP^{high}$ cells in each of these cell types. N = 3, n ≥ 7 embryos, and data represent mean ± SEM.

(B) Graphs show percentage chimerism (% $GFP^+$ cells) in granulocytes, lymphocytes and monocytes in peripheral blood at 1,2,3 and 4 months post HSC transplantation. N = 11 recipients/condition, data represent mean ± SEM.

(C) Graphs show percentage chimerism (% $GFP^+$ cells) in $CD4^+$ T cells, $CD8^+$ T cells, $CD19^+$ B cells, $CD11b^+$ myeloid cells, and $CD45^+$ pan hematopoietic cells in bone marrow harvested 4 months post HSC transplantation. N = 11 recipients/condition, data represent mean ± SEM.

(D) Representative flow cytometry scatter plot and graph show the frequency of MMP subsets in FL ST-HSCs. N = 5, n ≥ 5 fetal livers, data represent mean ± SEM.

(E) Graphs show MFI of EGFP in R26-mito-EGFP E15.5 fetal liver HSPC compartments. N = 5, n ≥ 5 fetal livers, and data represent mean ± SEM.

(F) Graphs show MFI of TMRM in WT E15.5 fetal liver HSPC compartments. N = 5, n ≥ 5 fetal livers, and data represent mean ± SEM.

**Fig S6. Single-cell RNA sequencing of E11.5 AGM MMP subsets.**

(A) Feature plots show expression of known AGM cell type markers- *Vegfc* (arterial endothelial cells), *Nrp2* (venous endothelial cells), *Cdh5* (Pre-HE cells), *Cldn5* (HE cells), *Plagl1* (Non-HE cells), *Col3a1* (mesenchymal cells), *Cdkn1c* (IAHCs), *Kit* (EHT cells), *Ikzf2* (Pro-HSCs), *Itgb3* (Pre-HSC type-I), *Csf1r* (Pre-HSC type-II). UMAP shows pooled datasets from two replicates. N = 2 independent experiments.

(B) Pie chart shows the distribution of WT E11.5 AGM cell types within the MMP$^{low}$ and MMP$^{high}$ subsets.

(C) Ridge plots show cell cycle analysis of MMP$^{low}$ and MMP$^{high}$ type-II HSCs based on expression levels cell cycle markers- *Pcna, Top2a, Mcm6* and *Mki67.*

(D) Dot plot shows expression levels of genes involved in mitochondrial quality control in MMP$^{low}$ and MMP$^{high}$ type-II HSCs.

(E) Cell Radar plots show blood cell lineage bias of pre-HSC type-II clusters.

(F) Dot plot shows the expression level of *Ociad1/asrij* in MMP$^{low}$ and MMP$^{high}$ type-II HSCs.

**Figure S7. Differential gene expression between MMP subsets of type-II HSCs.**

(A) Principal component analysis (PCA) of MMP$^{low}$ and MMP$^{high}$ type-II HSCs. N = 2.

(B) MA plot of MMP$^{low}$ and MMP$^{high}$ type-II HSCs.

(C) Heatmap shows expression levels of genes involved in the PI3K signaling axis in the MMP$^{low}$ and MMP$^{high}$ type-II HSCs. See table S6 for normalised gene counts.

(D) Dotplot shows the suppressed biological process in MMP$^{high}$ HSCs compared to MMP$^{low}$ HSCs.

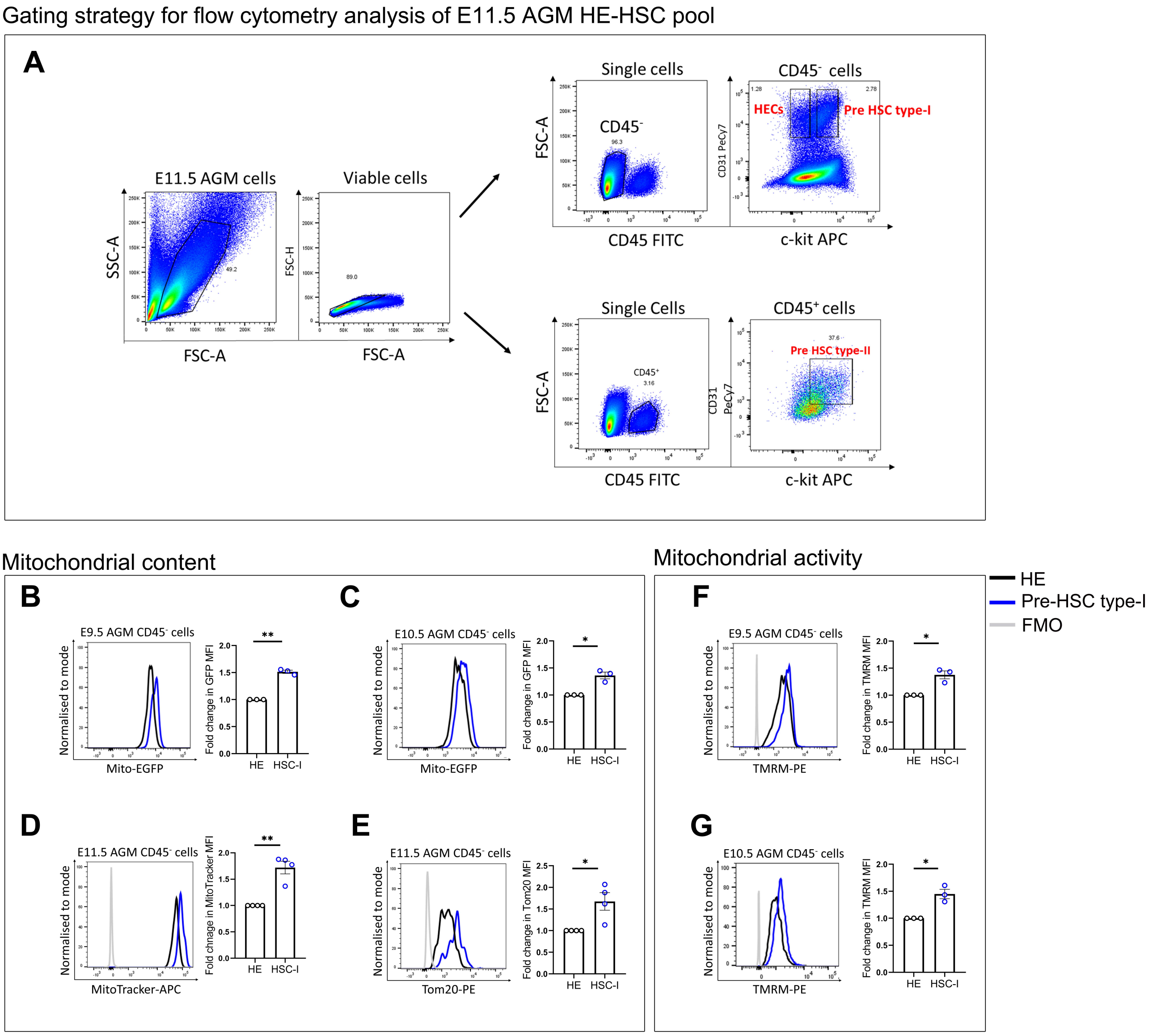

Gating strategy for flow cytometry analysis of E11.5 AGM HE-HSC pool
A
E11.5 AGM cells
SSC-A
FSC-A
49.2
Viable cells
FSC-H
FSC-A
89.0
Single cells
FSC-A
CD45-
96.3
CD45 FITC
CD45- cells
CD31 PeCy7
HECs
Pre HSC type-I
1.28
2.78
c-kit APC
Single Cells
FSC-A
CD45+
3.16
CD45 FITC
CD45+ cells
CD31 PeCy7
Pre HSC type-II
37.6
c-kit APC
Mitochondrial content
Mitochondrial activity
HE
Pre-HSC type-I
FMO
B
E9.5 AGM CD45- cells
Normalised to mode
Mito-EGFP
Fold change in GFP MFI
**
HE
HSC-I
C
E10.5 AGM CD45- cells
Normalised to mode
Mito-EGFP
Fold change in GFP MFI
*
HE
HSC-I
F
E9.5 AGM CD45- cells
Normalised to mode
TMRM-PE
Fold change in TMRM MFI
*
HE
HSC-I
D
E11.5 AGM CD45- cells
Normalised to mode
MitoTracker-APC
Fold chnage in MitoTracker MFI
**
HE
HSC-I
E
E11.5 AGM CD45- cells
Normalised to mode
Tom20-PE
Fold change in Tom20 MFI
*
HE
HSC-I
G
E10.5 AGM CD45- cells
Normalised to mode
TMRM-PE
Fold change in TMRM MFI
*
HE
HSC-I


**Figure S1**

Expression of mitochondrial genes in ckit$^{+}$ GATA2$^{low}$ HSC cluster

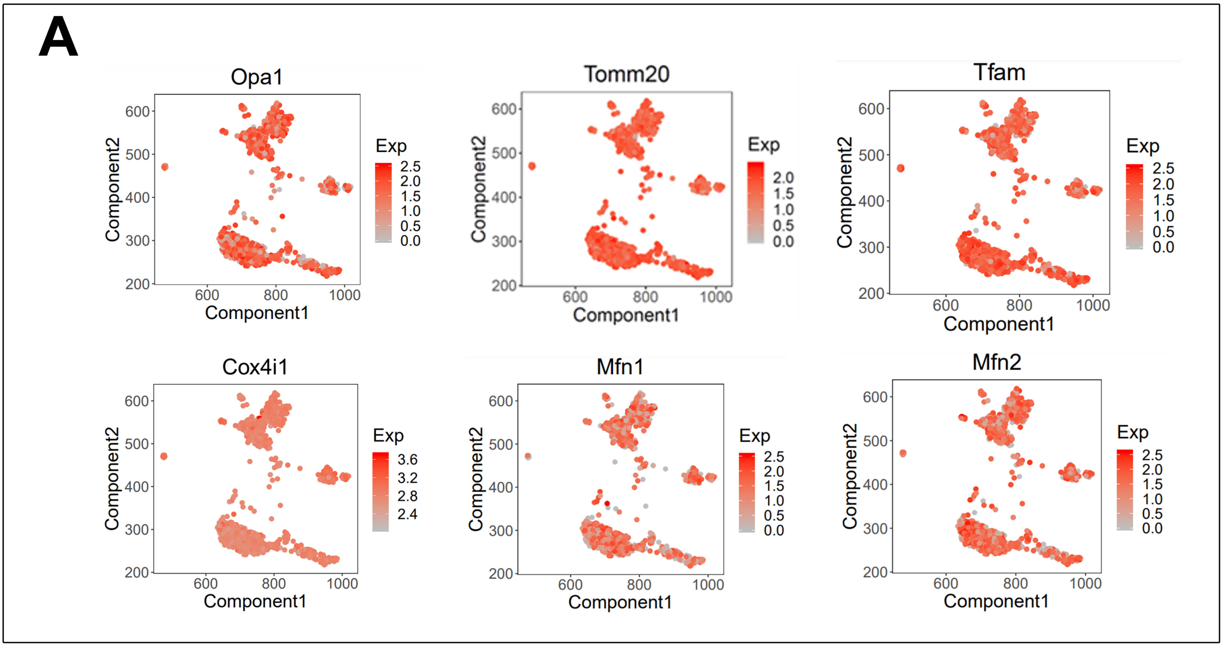


TMRM intensity profile in AGM cells

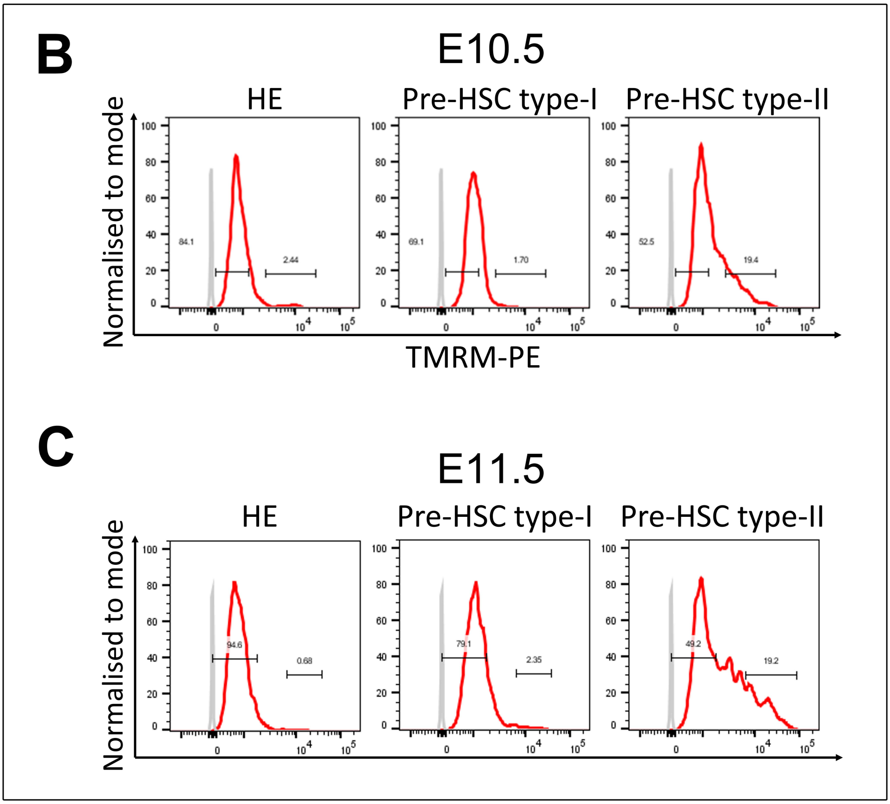


**Figure S2**

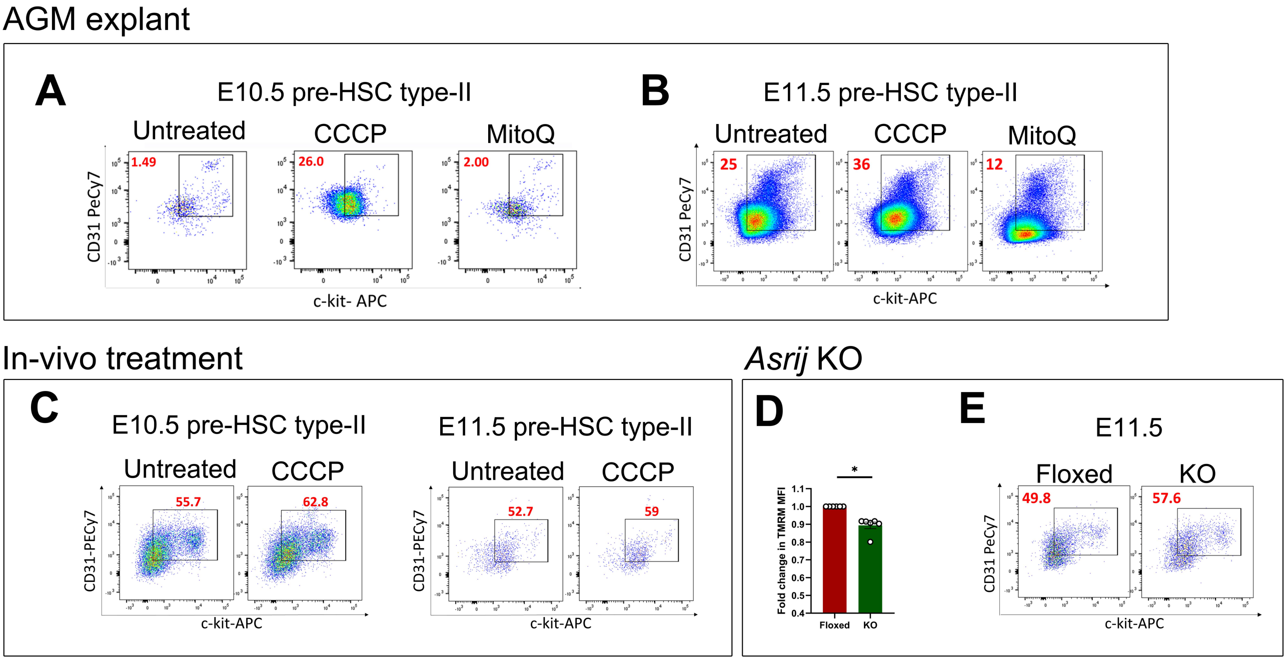
AGM explant
A
E10.5 pre-HSC type-II
Untreated
CCCP
MitoQ
1.49
26.0
2.00
CD31 PeCy7
c-kit- APC
B
E11.5 pre-HSC type-II
Untreated
CCCP
MitoQ
25
36
12
CD31 PeCy7
c-kit-APC
In-vivo treatment
C
E10.5 pre-HSC type-II
Untreated
CCCP
55.7
62.8
CD31-PECy7
c-kit-APC
E11.5 pre-HSC type-II
Untreated
CCCP
52.7
59
CD31-PECy7
c-kit-APC
Asrij KO
D
*
Fold change in TMRM MFI
Floxed
KO
E
E11.5
Floxed
KO
49.8
57.6
CD31 PeCy7
c-kit-APC

### Figure S3

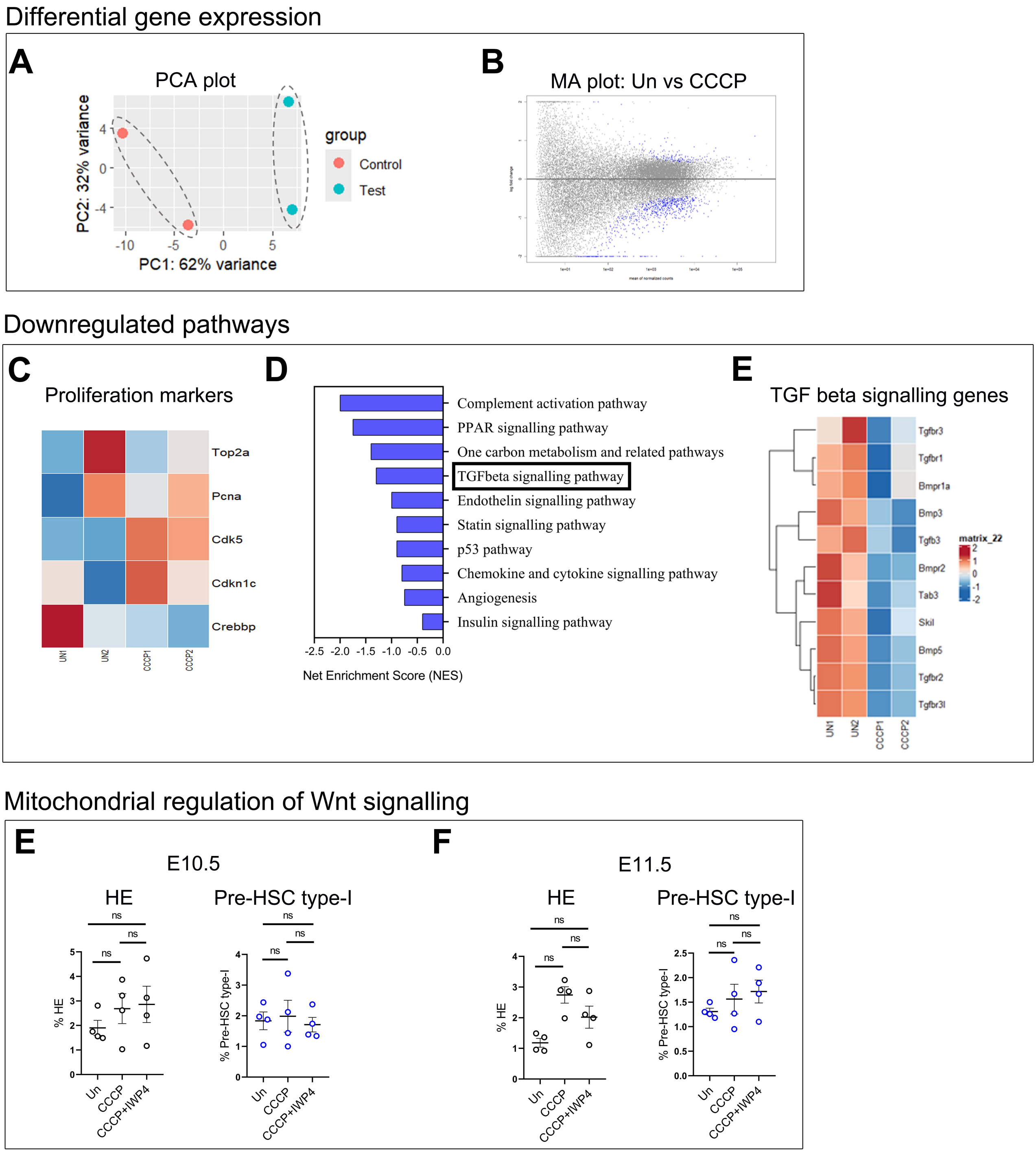
Differential gene expression
A
PCA plot
PC2: 32% variance
PC1: 62% variance
group
Control
Test
B
MA plot: Un vs CCCP
Downregulated pathways
C
Proliferation markers
Top2a
Pcna
Cdk5
Cdkn1c
Crebbp
UN1
UN2
CCCP1
CCCP2
D
Complement activation pathway
PPAR signalling pathway
One carbon metabolism and related pathways
TGFbeta signalling pathway
Endothelin signalling pathway
Statin signalling pathway
p53 pathway
Chemokine and cytokine signalling pathway
Angiogenesis
Insulin signalling pathway
Net Enrichment Score (NES)
E
TGF beta signalling genes
Tgfbr3
Tgfbr1
Bmpr1a
Bmp3
Tgfb3
Bmpr2
Tab3
Skil
Bmp5
Tgfbr2
Tgfbr3l
matrix_22
Mitochondrial regulation of Wnt signalling
E
E10.5
HE
Pre-HSC type-I
% HE
% Pre-HSC type-I
ns
Un
CCCP
CCCP+IWP4
F
E11.5
HE
Pre-HSC type-I
% HE
% Pre-HSC type-I
ns
Un
CCCP
CCCP+IWP4

**Figure S4**

#### MMP based distribution of E10.5 AGM cells

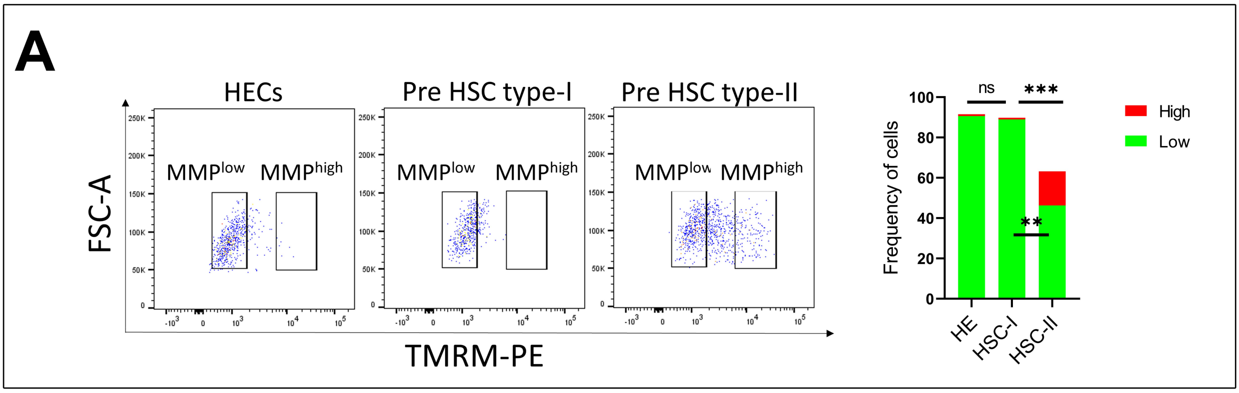


#### Peripheral blood and BM chimerism

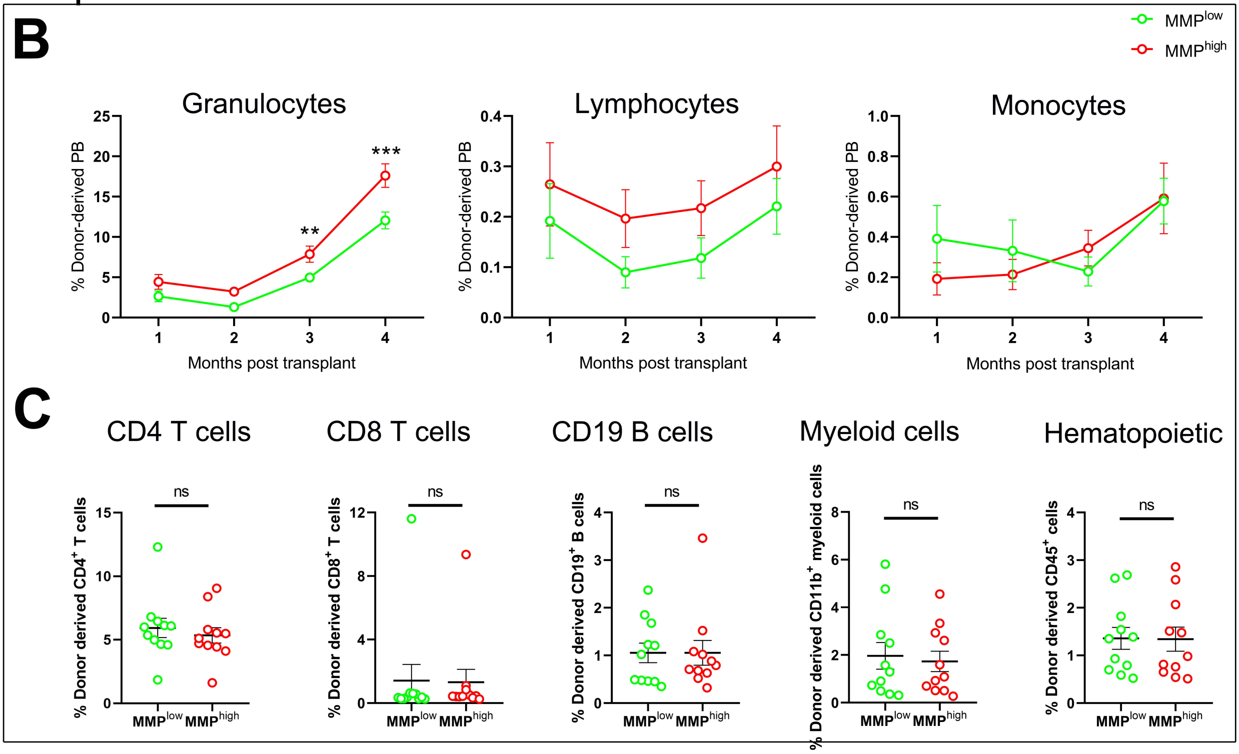


#### E15.5 fetal liver

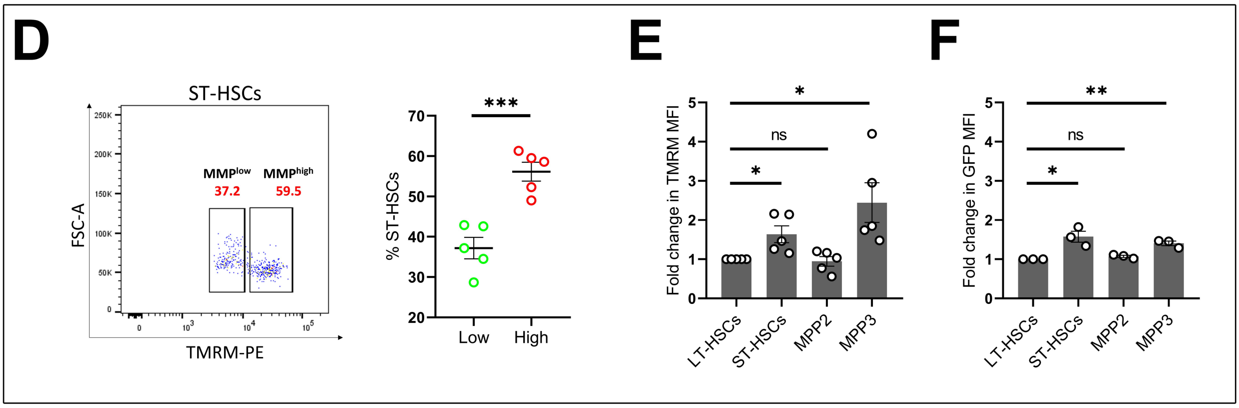


**Figure S5**

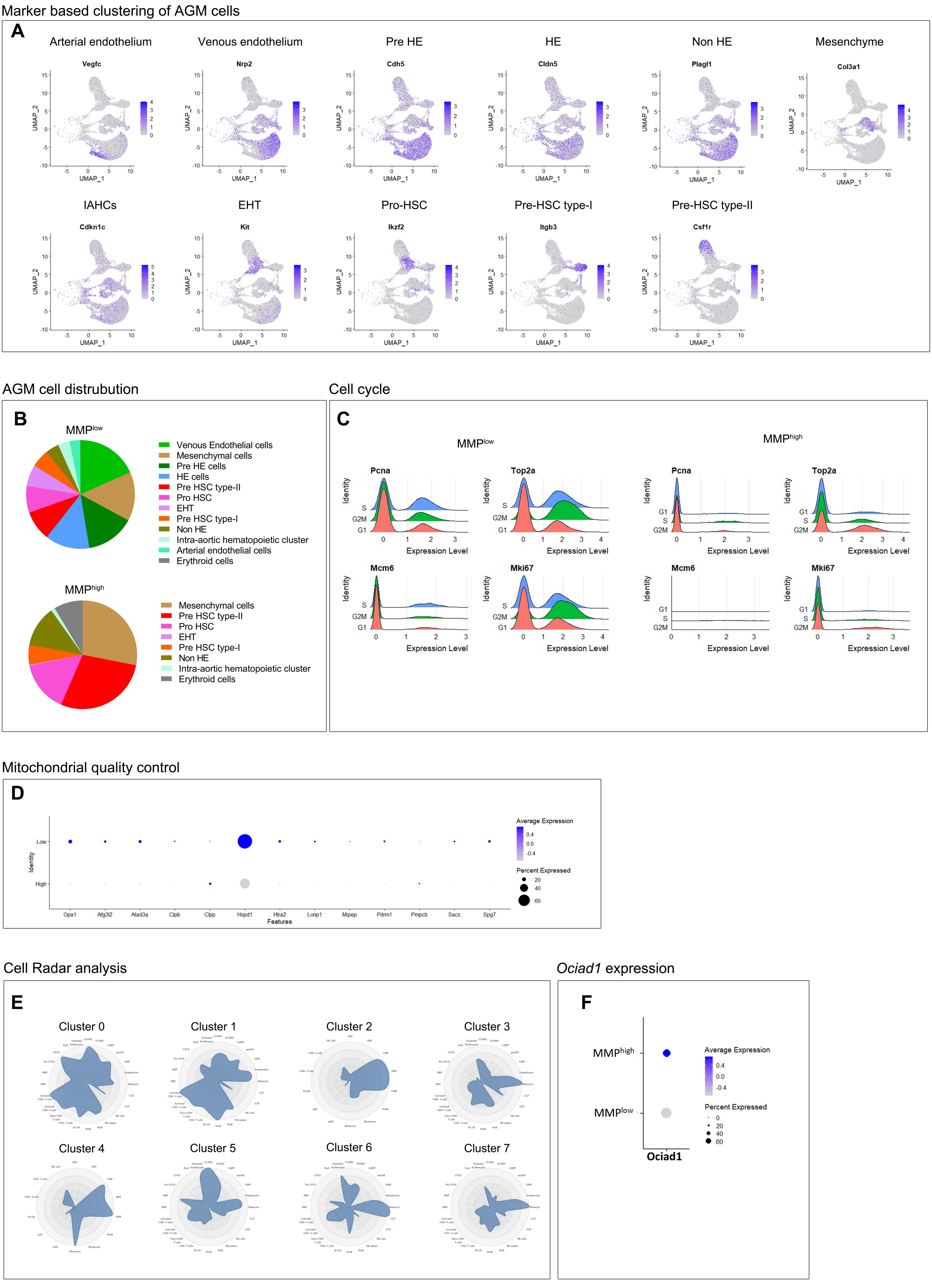
Marker based clustering of AGM cells
A
Arterial endothelium
Vegfc
Venous endothelium
Nrp2
Pre HE
Cdh5
HE
Cldn5
Non HE
Plagl1
Mesenchyme
Col3a1
IAHCs
Cdkn1c
EHT
Kit
Pro-HSC
Ikzf2
Pre-HSC type-I
Itgb3
Pre-HSC type-II
Csf1r
UMAP_1
UMAP_2
AGM cell distrubution
B
MMPlow
Venous Endothelial cells
Mesenchymal cells
Pre HE cells
HE cells
Pre HSC type-II
Pro HSC
EHT
Pre HSC type-I
Non HE
Intra-aortic hematopoietic cluster
Arterial endothelial cells
Erythroid cells
MMPhigh
Mesenchymal cells
Pre HSC type-II
Pro HSC
EHT
Pre HSC type-I
Non HE
Intra-aortic hematopoietic cluster
Erythroid cells
Cell cycle
C
MMPlow
MMPhigh
Pcna
Top2a
Mcm6
Mki67
Identity
S
G2M
G1
Expression Level
Mitochondrial quality control
D
Low
High
Identity
Opa1
Afg3l2
Atad3a
Clpb
Clpp
Hspd1
Htra2
Lonp1
Mipep
Pitrm1
Pmpcb
Sacs
Spg7
Features
Average Expression
Percent Expressed
Cell Radar analysis
E
Cluster 0
Cluster 1
Cluster 2
Cluster 3
Cluster 4
Cluster 5
Cluster 6
Cluster 7
Ociad1 expression
F
MMPhigh
MMPlow
Ociad1
Average Expression
Percent Expressed

**Figure S6**

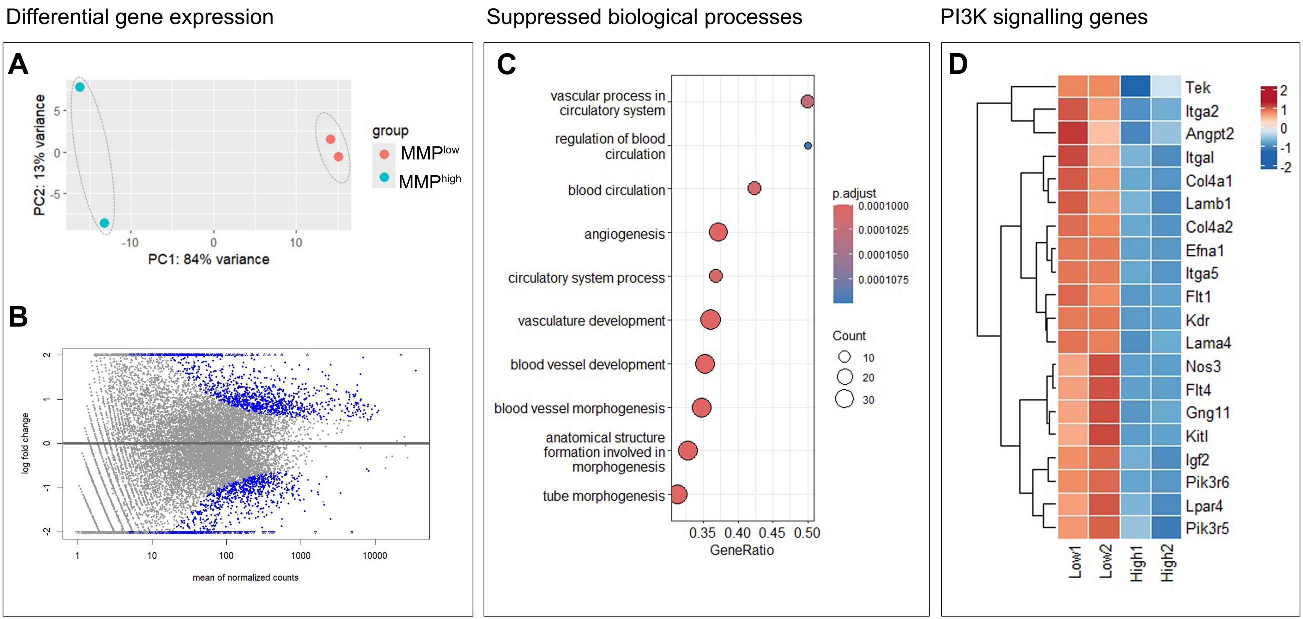
Differential gene expression
Suppressed biological processes
PI3K signalling genes
A
PC2: 13% variance
PC1: 84% variance
group
MMPlow
MMPhigh
B
log fold change
mean of normalized counts
C
vascular process in circulatory system
regulation of blood circulation
blood circulation
angiogenesis
circulatory system process
vasculature development
blood vessel development
blood vessel morphogenesis
anatomical structure formation involved in morphogenesis
tube morphogenesis
GeneRatio
p.adjust
0.0001000
0.0001025
0.0001050
0.0001075
Count
10
20
30
D
Tek
Itga2
Angpt2
Itgal
Col4a1
Lamb1
Col4a2
Efna1
Itga5
Flt1
Kdr
Lama4
Nos3
Flt4
Gng11
Kitl
Igf2
Pik3r6
Lpar4
Pik3r5
Low1
Low2
High1
High2

**Figure S7**

**Table S1. Primer sequences**

| S.No | Gene | Sequence |
|---|---|---|
| 1 | *Tfam* | Forward: 5'-GTCCATAGGCACCGTATTGC-3'<br>Reverse: 5'-CCCATGCTGGAAAAACACTT-3' |
| 2 | *Idh* | Forward 5'-AATGCTCAAGCCAACTCTCC-3'<br>Reverse 5'-TTGTTTGTTGTGAGGAAATGC-3 |
| 3 | *Pgc1α* | Forward 5'-GTAGGCCCAGGTACGACAGC-3'<br>Reverse 5'-GCTCTTTGCGGTATTCATCCC-3' |
| 4 | *Atp5a* | Forward 5'-TGGAAACAATGTATTCGTTCAT-3'<br>Reverse 5'-CTCCTCCGGACGCAAAT-3' |
| 5 | *Nrf1* | Forward 5'-CAACAGGGAAGAAACGGAAA-3'<br>Reverse 5'-GCACCACATTCTCCAAAGGT-3' |
| 6 | *Nrf2* | Forward 5'-AGGTTGCCCACATTCCCAAACAAG-3'<br>Reverse 5'-TTGCTCCATGTCCTGCTCTATGCT-3' |
| 7 | *β2M* | Forward: 5'-ACAGTTCCACCCGCCTCACATT-3'<br>Reverse: 5'-TAGAAAGACCAGTCCTTGCTGAAG-3' |
| 8 | *mt-Cytb* | Forward: 5'-ACACGCAAACGGAGCCTCAA-3'<br>Reverse: 5'-TGCTGTGGCTATGACTGCGAACA-3' |
| 9 | *mt-Nd2* | Forward: 5'-CCTCCTGGCCATCGTACTCA-3'<br>Reverse: 5'-GAATGGGGCGAGGCCTAGTT-3' |
| 10 | *Sdh* | Forward: 5'-GCTCGAGCTCTCCTACTCC-3'<br>Reverse: 5'-GCTTGGTGACAGGTGAATGT-3' |
| 11 | *Hk2* | Forward: 5'-ATTGTGGCTGTGGTGAA-3'<br>Reverse: 5'-AATGTGACGCATCTCCTC-3' |
| 12 | *Pfkfb3* | Forward: 5'-AGCCTCTTGACCCTGATA-3'<br>Reverse: 5'-TTCTTGCCTCTGCTGGAC-3' |
| 13 | *Hif1α* | Forward: 5'-CCTGCACTGAATCAAGAGGTTGC-3'<br>Reverse: 5'-CCATCAGAAGGACTTGCTGGCT-3' |
| 14 | *PGK cre* | Forward: 5'- TTC TCC TCT TCC TCA TCT CCG-3'<br>Reverse: 5'-GCA AAC GGA CAG AAG CAT TT-3' |
| 15 | *Mito-EGFP* | Forward: 5'-TCCCTCGTGATCTGCAACTCCAGTC-3'<br>Reverse: 5'-AACCCCAGATGACTACCTATCCTCC-3' |
| 16 | *Asrij floxed* | Forward: 5'-GGAGAATTGCGGCGCTCTTCTCC -3'<br>Reverse: 5'- CCATCCATCCCTCTCCACTGG -3' |
| 17 | *Asrij KO* | Forward: 5'-ATGAAGCAGTGTCTTGGGATTGC-3'<br>Reverse: 5'-CCATCCATCCCTCTCCACTGG-3' |
